# Costs of choosiness can promote reproductive isolation in parapatry

**DOI:** 10.1101/2020.11.12.379461

**Authors:** Thomas G. Aubier, Mathieu Joron

## Abstract

Species often replace each other spatially along contact zones, giving scope for parapatric speciation. In models of parapatric speciation driven by assortative mating, costs of female choosiness have so far be ignored. Yet, if females encounter only a limited number of males, those that are too choosy may remain unmated, and this should create direct sexual selection against choosiness. In our individual-based model of parapatric ecological speciation, disruptive viability selection leads to divergence of an ecological trait. Additionally, female choosiness (following a ‘matching mating rule’ based on the same ecological trait) can evolve at the risk of remaining unmated, and can limit gene flow between diverging populations. In line with previous litterature, out of the contact zone, the evolution of cost-free choosiness stops at intermediate values due to indirect selection against strong choosiness. Here we show that a weak cost of choosiness, by modifying genotypic frequencies on which viability selection acts, reduces this selection pressure, thus permitting the evolution of stronger choosiness than in the absence of costs. In strong contrast to sympatric models, costs of choosiness can therefore promote reproductive isolation in parapatry.

## INTRODUCTION

Since the publication of Darwin’s (1859) *The Origin of Species*, a large amount of research has been carried out on speciation (see Turelli et al., 2001; The Marie Curie Speciation Network, 2012; White et al., 2020, for a review). Traditionally, speciation events have been categorized according to their geographical context – allopatry (extrinsic barrier to genetic exchange during divergence), parapatry (partial extrinsic barrier) and sympatry (no extrinsic barrier). The allopatric mode of speciation had been viewed as the most plausible one for decades, until mounting empirical evidence suggested that speciation with gene flow (i.e., in sympatry or parapatry) can occur and might be common in nature (Rice and Hostert, 1993; Via, 1999; Barluenga et al., 2006; Savolainen et al., 2006; Soria-Carrasco et al., 2014; Seehausen et al., 2014; Momigliano et al., 2017). Mathematical models have since been developed to dissect the factors involved in speciation with gene flow (see Gavrilets, 2003, 2014; Kopp et al., 2018). From this point on, research on speciation has shifted from a focus on the geographical context of differentiation to a focus on the role of viability and sexual selection (through differential viability and mating success among genotypes, respectively) in the evolution of reproductive isolation (Via, 2001; Kirkpatrick and Ravigné, 2002; Kopp et al., 2018) and in the maintenance of gene flow that is pervasive in nearly all systems (Mallet, 2005; Nosil et al., 2009; Marques et al., 2019). It is now widely accepted that geography is but one factor among others that can influence the progress toward speciation.

Viability and sexual selection can act on heritable traits if individuals differ in their survival rate and mating success, respectively. In particular, disruptive selection – when individuals with intermediate phenotypic traits have a low fitness – plays a key role for population divergence and speciation (Kirkpatrick and Ravigné, 2002). First, abiotic or biotic environmental factors can cause disruptive viability selection if individuals with intermediate phenotypes suffer viability costs (e.g., high predation risk, low efficiency in resource use, genetic incompatibilities). Second, under the right circumstances, mate preferences can cause disruptive sexual selection on traits used as phenotypic cues for mate choice. In particular, if individuals express a preference for mates of their own phenotype (‘matching mating rule’, described below), the mating success associated with a phenotype increases with its frequency (positive frequency-dependent sexual selection). Therefore, individuals with intermediate phenotypes may have a low mating success if they are rare; sexual selection may be disruptive (Kirkpatrick and Nuismer, 2004; Pennings et al., 2008; Otto et al., 2008). Such disruptive selection – either via reduced survival or via reduced mating success of individuals with intermediate phenotypes – strongly contributes to the divergence of populations with distinct adaptive phenotypes (Kirkpatrick and Ravigné, 2002).

Assortative mating – i.e., the tendency of individuals of similar phenotype to mate together more often than expected at random – plays a key role in generating reproductive isolation. It can arise as a by-product of adaptive divergence via temporal or spatial isolation (e.g., mediated by the evolution of host choice or phenology; Servedio et al., 2011), or can be driven by various behavioural processes (Kirkpatrick and Ravigné, 2002; Cézilly, 2004). Of particular interest is the case of matching mating rules (Kopp et al., 2018), where individuals preferentially choose mates with which they share phenotypic traits such as colours (Summers et al., 1999; Jiggins et al., 2001; Reynolds and Fitzpatrick, 2007; Bortolotti et al., 2008) or acoustic signals (Snowberg and Benkman, 2007). Disruptive selection can indirectly favour the evolution of choosiness (hereafter, choosiness refers to the strength of preferences for ‘matching’ mates) through linkage disequilibrium (when alleles for strong choosiness are correlated with alleles coding for extreme ecological phenotypes), thereby leading to premating isolation between diverging populations (Rundle and Nosil, 2005; Sobel et al., 2010). This process can be facilitated if mate choice is directly based on the traits under disruptive selection (so-called ‘magic traits’; Servedio et al., 2011).

This recent focus on the role of viability and sexual selection in reproductive isolation actually put the geographical context aside, especially from a theoretical perspective. Gavrilets (2014) noted that parapatric speciation is the least theoretically studied geographical mode of speciation, whereas he argued that it should be “the most general” one (Gavrilets, 2003). Environmental heterogeneity taking the form of ecological gradients is pervasive in natural systems, and applies to both abiotic factors (e.g., temperature, humidity, altitude) and biotic factors (e.g., predation risk). While ecological gradients setting the stage for parapatric speciation are ubiquitous, most theoretical studies have focused on sympatric speciation, maybe because speciation in parapatry seems more straightforward to understand than sympatric speciation (Dieckmann and Doebeli, 1999; Matessi et al., 2001; Bolnick, 2004; Bürger et al., 2006; Kopp and Hermisson, 2008; Otto et al., 2008; Thibert-Plante and Hendry, 2011; Rettelbach et al., 2011; Aubier et al., 2019). Indeed, limited dispersal and reduced survival of immigrants in foreign environments can be seen as just an additional isolating barrier (Kirkpatrick and Ravigné, 2002; Nosil et al., 2005). Nevertheless, spatial structure has been shown to facilitate speciation – potentially even more than complete spatial isolation – and the ecological contact is actually the driving force leading to reproductive isolation in parapatry (Doebeli and Dieckmann, 2003).

Along ecological gradients, diverging populations replace each other spatially. The relative proportions of phenotypes change and can be unbalanced as one moves away from the centre of the gradient, with each phenotype being predominant at one of the two edges of the gradient. This property can strongly affect density- and frequency-dependent forces acting during speciation and can lead to outcomes that are specific to the parapatric context. In particular, positive frequency-dependent sexual selection caused by female choosiness can promote ecological divergence by favouring different phenotypic traits in parapatry (Servedio, 2011). However, in such parapatric context, very strong female choosiness leads rare males to mate with rare (matching) females, resulting in equal mating success among males and therefore in the loss of sexual selection. Such loss of sexual selection under strong choosiness causes a positive genetic association between choosiness and locally *disfavoured* ecological alleles. This ultimately leads to the removal of alleles coding for strong choosiness by indirect viability and sexual selection. As a result, only intermediate choosiness can evolve in parapatry (Servedio, 2011; Cotto and Servedio, 2017, see also Appendix A). Note that the situation is rather different when the build-up of assortative mating is based on a different genetic architecture; Servedio and Burger (2014) showed that sexual selection inhibits differentiation when mate choise is based on a preference/trait rule.

The distribution of diverging populations in parapatry may generate variation in mating success among females. In particular, variation in female mating success can take the form of a cost of choosiness – i.e., a fitness cost to females that are choosy during mate choice –, thereby generating direct sexual selection inhibiting choosiness. Choosy females that refuse to mate with unpreferred males are more likely to delay mating than nonchoosy ones, possibly generating variation in mating success among females. When females may encounter a limited number of males in their lifetime, delaying mating may even lead choosy females to remain unmated (Reynolds and Gross, 1990; Milinski et al., 1992; Heubel et al., 2008; Scott et al., 2020) (but see Puebla et al., 2012), especially under limited energetic reserves (Byers et al., 2005), high predation risk (Bonachea and Ryan, 2011), or low population density. Such direct sexual selection against choosiness can inhibit the evolution of choosiness in sympatry (Bolnick, 2004; Schneider and Bürger, 2006; Bürger et al., 2006; Kopp and Hermisson, 2008). In many theoretical models, however, female mating probabilities are normalized, so that all females eventually find a mate (Dieckmann and Doebeli, 1999; Doebeli and Dieckmann, 2000, 2003; Servedio, 2011; Thibert-Plante and Gavrilets, 2013), while this sexual selection pressure acting on choosiness is likely to depend on the local distribution of the phenotypes used as bases of mate choice. If the frequencies of the diverging populations are balanced, females often, if not always, end up mating with one of the males they prefer as long as they encounter enough males in their lifetime. Thus, when modelling speciation in sympatry, theoreticians can assume that all females eventually find a mate without much loss of generality. In parapatry, however, females with rare phenotypes (i.e., immigrant females) strongly delay mating, and risk to remain unmated if they are too choosy, even if they can encounter many males in their lifetime. Yet, so far, this risk has been ignored when modelling parapatric speciation (Doebeli and Dieckmann, 2003; Kawata et al., 2007; Leimar et al., 2008; Ispolatov and Doebeli, 2009; Thibert-Plante and Gavrilets, 2013; Cotto and Servedio, 2017) (note that M’Gonigle et al., 2012, modelled this risk in a parapatric spatial setting but have not considered its implication for the evolution of choosiness). Whether such direct sexual selection against choosiness inhibits the build-up of reproductive isolation in parapatry remains to be investigated.

With stochastic individual-based simulations, we here test if the risk of remaining unmated incurred by choosy females (hereafter called ‘cost of choosiness’ because only choosy females reject potential mates at the risk of remaining unmated) inhibits the evolution of choosiness in parapatry. We consider that two ecotypes are favoured in distinct habitats (i.e., with distinct abiotic or biotic environment) in parapatry. We implement two ‘main demes’ where viability selection is directional (with divergent optima between demes), and a ‘transition deme’ where viability selection is disruptive (both ecotypes are favoured and hybrid individuals suffer viability costs). Such spatial discretization allows us to infer the global selection gradient from local selection gradients (using a method derived from Wickman et al., 2017). The ecological trait under disruptive viability selection can be used as the basis of mate choice by choosy females (matching mating rule), and the evolution of female choosiness can therefore lead to assortative mating and reproductive isolation between ecotypes. By assuming females may only encounter a limited number of males in their lifetime, we account for the possibility that choosy females may risk to remain unmated, in particular if matching males are locally rare, thereby generating variation in mating successes among females. Note that for simplicity, we assume that females do not adjust their level of choosiness in response to the abundance of males from their own type (as shown by Willis et al., 2011). In line with previous litterature, we show that in the main demes, the evolution of cost-free choosiness stops at intermediate values due to indirect viability selection against strong choosiness. We also confirm that a cost of choosiness generates additional sexual selection pressure against choosiness. But in strong contrast to sympatric models, interestingly, we find that a weak cost of choosiness can favour the evolution of choosiness by impairing the build-up of allelic associations that would cause indirect viability selection against strong choosiness. As a result, our model predicts that the effects of the cost of choosiness on reproductive isolation in parapatry greatly depend on the spatial structure and on the strength of viability selection.

## METHODS

### Individual characteristics and spatial context

Individuals are characterized by their sex, their ecological trait *x* ∈ [0,1] controlling survival in a local environment, and their choosiness trait *c* ∈ ℝ controlling the strength of female preference during mate choice. The genetic architecture and inheritance rules are based on Claessen et al. (2008). The traits x and c are respectively determined by *L_x_* and *L_c_* diploid unlinked loci on autosomal chromosomes. Each allele can take any real value in the range of the trait it is coding for – i.e., in [0,1] and ℝ for the alleles coding for traits *x* and *c*, respectively. Trait *x* and *c* are calculated as the mean values of their alleles. In addition, to measure genetic differentiation at neutral loci caused by assortative mating, we assume that individuals carry *L_n_* diploid unlinked neutral loci with a large number (= 50) of possible alleles (as in Thibert-Plante and Hendry, 2009; Irwin, 2020).

Individuals are distributed in two main demes with distinct directional viability selection pressures (subscript 1 and 2) and in a transition deme with a disruptive viability selection pressure (subscript T), with carrying capacities *K*_1_, *K*_2_ and *K*_T_ respectively (see the flowchart of the model, Fig. 1). The overall carrying capacity *K* is set to be constant (*K* = *K*_1_ + *K*_2_ + *K*_T_; such that the extent of genetic drift does not vary too much across simulations) and we assume that demes 1 and 2 have the same carrying capacity (*K*_1_ = *K*_2_). The carrying capacity of each deme depends on the relative size of the transition deme, called *α*_T_(*α*_T_ ∈ [0,1]). In particular, the carrying capacity of the transition deme is calculated as: *K*_T_ = *α*_T_*K*.

**Figure 1:**
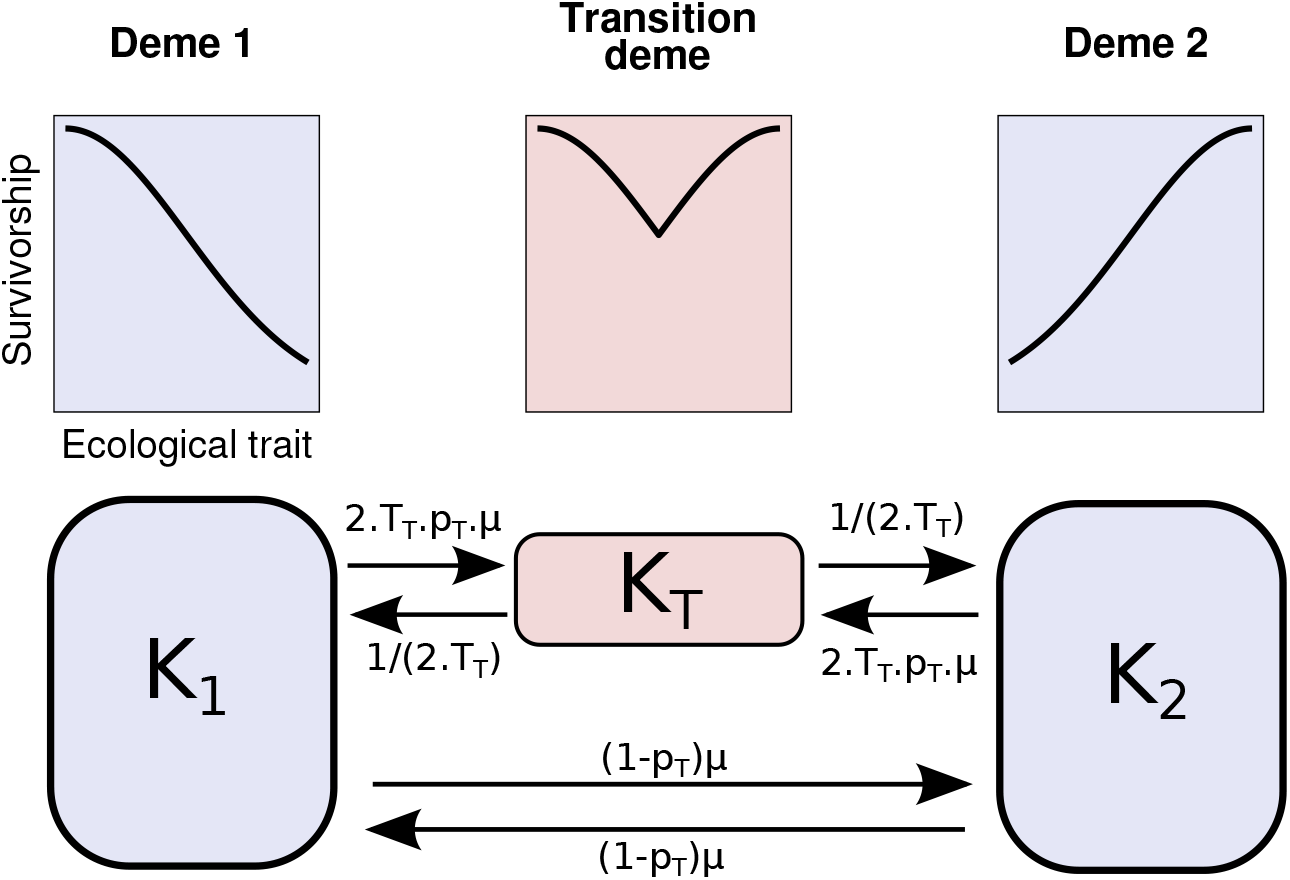
Flowchart of the model: individual survivorship, carrying capacities and migration rates. The main demes and the transition deme are coloured in blue and pink, respectively.

During their lifetime, individuals may experience three processes in this given order: survival to adult state, dispersion and reproduction. We assume that generations do not overlap.

### Survival to adult state

Viability costs, which affect survival to adult state, are heterogeneous in space. Through differential survivorship among individuals, viability selection acts on the ecological trait (Fig. 1). In main demes, viability selection is directional. The ‘condition’ *ω_i_* of one individual in a main deme *i* ∈ {1,2} depends on its ecological trait *x*. In particular, if *x* ≃ *θ_i_*, then the individual does not incur viability costs:

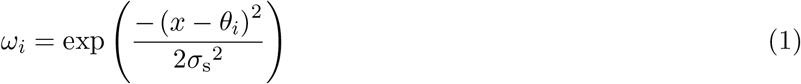

Where *θ_i_* is the local optimal trait value and *σ_s_* is inversely proportional to the strength of viability selection, which is the same in both demes. We implement *θ*_1_ = 0 and *θ*_2_ = 1. Therefore, individuals with high viability in deme 1 suffer from low viability in deme 2, and vice versa.

In the transition deme, viability selection is disruptive in a way that individuals exhibiting ecological traits matching the optimal trait values *θ*_1_ or *θ*_2_ do not incur viability costs. On the contrary, individuals with an intermediate ecological trait suffer from high viability costs:

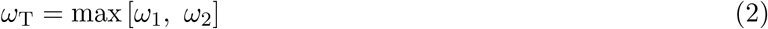

Thus, we assume that *σ_s_* is the same in the three demes.

In addition to individuals’ condition, density-dependent competition for resources will define individuals’ survival to adult state. To do so, we use the Beverton-Holt equation – i.e., the discrete version of the logistic growth – to calculate the survival probability *v* to adult state in deme *i* ∈ {1, 2, T}:

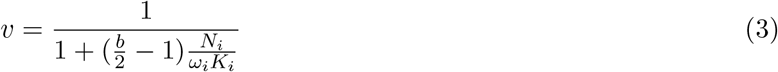

With *b* the mean number of offspring per female, *N_i_* the number of individuals in deme *i* and *K_i_* the carrying capacity of the population in deme *i*.

### Dispersal

Once individuals reach their adult state, they can migrate to other demes, before the reproduction stage.

We implement *μ* as the overall migration rate between the two main demes. This parameter affects all migration flows, and therefore represents the dispersal ability. Depending on the size of the transition deme, a fraction *p*_T_ of migrant lineages will experience the transition deme during a number of generations *T*_T_ before migrating again – i.e., a portion 1/*T*_T_ of individuals from the transition deme are migrating at each generation. Another fraction 1 – *p*_T_ will migrate directly to the other main deme, without experiencing the transition deme. We can then calculate all migration flows from the parameters *μ*, *p*_T_ and *T*_T_ (see the flowchart of the model, Fig. 1).

The relative size of the transition deme (*α*_T_) will influence the intensity of these migration flows. A relationship between *p*_T_, *T*_T_, *μ* and *α*_T_ can be calculated if we apply a reciprocity rule to the number of individuals migrating from one deme to another. If we consider that there is the same number of migrants transiting from one main deme (say deme 1) to the transition deme than vice-versa, we get:

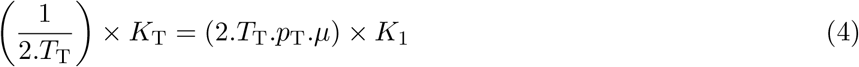

Together with the relationship *K*_T_ = *α*_T_.*K*, we can calculate:

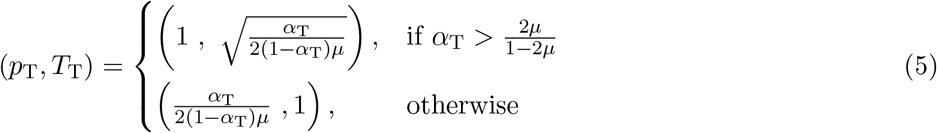

Therefore, with a large transition deme (high *α*_T_), lineages of migrants leaving the main demes (*p*_T_ = 1) experience on average *T*_T_ > 1 generations in the transition deme. With a small transition deme (low *α*_T_), only a fraction *p*_T_ < 1 of those migrants experiences on average one generation in the transition deme (*T*_T_ = 1).

### Reproduction

During the reproduction phase, each female is courted sequentially by a series of *N*_male_ males randomly chosen in her deme. We assume that males can mate more than once, do not exhibit any preferences and try to court all the females they encounter. On the contrary, we consider that a female can only mate once during her lifetime. Therefore, at each encounter, she evaluates the male and she decides to mate or not accordingly. If a choosy female does not choose to mate with any of the *N*_male_ males encountered, she does not mate at all. If *N*_male_ = ∞, the female keeps being presented with males (even those that have already been presented) until she does accept one; in that case, a very choosy female is likely to end up mating with a male she prefers even if those preferred males happen to be rare locally.

To evaluate a potential mate, a female is using the male ecological trait as the basis of choice. Depending on her choosiness trait c and on the difference between their ecological traits, the female will choose to mate with a probability Ψ following a ‘matching rule’ (Kopp et al., 2018). We use the preference function defined by Carvajal-Rodriguez and Rolán-Alvarez (2014) (Fig. S1). It satisfies basic properties – monotonicity, proportionality and symmetry –, which are necessary to avoid some modelling artifacts during the evolution of choosiness:

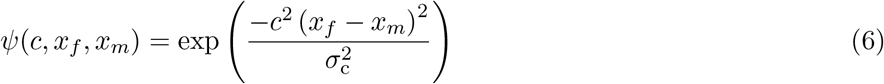

With *x_f_* and *x_m_* the ecological traits of the female and the male respectively, *c* the female choosiness trait and *σ_c_* the strength of the preference, which makes the mating probability more or less sensitive to *c*. Mating is random if *c* = 0 (*ψ* = 1; i.e., the female mate with the first male encountered), whereas it is positive assortative if *c* > 0. A female with choosiness *c* > 0 would mate preferentially with a male matching her own ecological trait – i.e., *c* > 0 indicates her readiness to reject males that do not match her ecological trait (*ψ* < 1).

After mating, the number of offspring is chosen randomly and independently from a Poisson distribution with parameter *b* – i.e., the average number of offspring per female. The sex of each offspring is chosen randomly assuming a balanced sex ratio. At the genetic level, we assume independent segregation. At each locus, one maternal allele and one paternal allele are randomly chosen. Mutation can occur at each allele determining trait *k* (*k* ∈ {*c, x*}) with a probability *m_k_*. The mutant allele value is drawn from a truncated normal distribution centered on the parent allele value with deviation 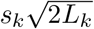 with 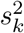 the mutational variance at the trait level and *L_k_* the number of loci determining the trait *k* (Aguilée et al., 2013), with allele values constrained to be in the interval [0, 1]. Mutation on each neutral locus occurs with probability *m_n_*, in which case a random allele is chosen from the 50 possible alleles.

As we assume non-overlapping generations, all adult individuals die after their reproduction phase, and the population is then entirely reconstituted from offspring.

### Numerical simulations and summary statistics

Our initial conditions correspond to a secondary contact scenario, where all individuals are specialists within their environment. All individuals have an ecological trait *x* = 0 in deme 1 and *x* = 1 in deme 2. In the transition deme, half of the population have an ecological trait *x* = 0 while the other half have *x* = 1. All individuals have a choosiness trait *c* = 0.2 (to insure that polymorphism is initially maintained). All neutral marker alleles are randomly picked, therefore there is no genetic differentiation at neutral loci among differentiated populations (while unrealistic, this assumption allows us to pinpoint the genetic differentiation caused by the evolution of choosiness).

For each simulation, we record the distribution of the ecological and choosiness traits, both in the main demes and in the transition deme. To assess the level of gene flow in our model, we measure the frequency of strong nonrandom mating and the genetic differentiation in neutral marker traits between ecotypes. We arbitarily define two ecotypes based on individuals’ ecological traits *x*; one ecotype is characterized by all individuals with *x* ≤ 0.5 and the other ecotype by all individuals with *x* > 0.5. Defining the different ecotypes as the populations from the two main demes (whatever the individuals’ ecological traits) leads to the same results (Fig. S2). To detect strong nonrandom mating caused by choosiness, we use a simple statistics 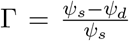 where *ψ_d_* and *ψ_s_* are the mean preference scores ψ between two individuals from different and the same ecotype respectively (see Equation 6 for the definition of the preference score; following Thibert-Plante and Gavrilets, 2013). Γ reflects the strength of reproductive isolation between ecotypes via nonrandom mating. Specifically, we consider that *strong nonrandom mating* has evolved if Γ > 0.99, so that the probability of mating between individuals from different ecotype is at least 100 times smaller than that between two individuals from the same ecotype (irrespective of their localisation). To assess the level of reproductive isolation among ecotypes, we compute the genetic differentiation *F*_ST_ statistics at neutral loci: 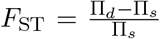. Π_*d*_ and Π_*s*_ represent the average number of pairwise differences in neutral loci between two individuals sampled from different ecotypes (Π_*d*_) or from the same ecotype (Π_*s*_). Therefore, in our numerical simulations, the *F*_ST_ index between ecotypes reflects the strength of reproductive isolation between ecotypes. We get qualitatelively the same result when considering the absolute genetic difference between ecotypes (Π_*d*_ – Π_*s*_, Fig. S3)

In numerical simulations, we vary the relative size of the transition deme (*α*_T_ = {0, 0.1, 0.2}), the strength of selection (*σ_s_* = {0.2, 0.3, 0.4, 0.5}) and the maximum number of males encountered per female (*N*_male_ = {1, 2, 4, 8, 16, 32, 64, ∞}). We constrain the range of *α*_T_ and *σ*_s_ analyzed to prevent the loss of polymorphism at the ecological locus (by genetic swamping; Lenormand, 2002) or the formation of a monomorphic transition deme (Fig. S4). The other parameters are fixed: *K* = 10, 000, *μ* = 0.01, *σ*_c_ = 0.2, *b* = 5, *L_x_* = 4, *L_c_* = 1, *m_x_* = *m_c_* = 5.10^-2^, *s_x_* = *s_c_* = 0.002, *L_n_* = 20, *m_n_* = 10^-4^. Note that we do not aim to perform an exhaustive study of the factors affecting the evolution of choosiness in parapatry (see more complete sensitivity analyses in Thibert-Plante and Gavrilets, 2013; Cotto and Servedio, 2017). We intentionally choose optimal values of parameters for choosiness to evolve without loss of polymorphism (secondary contact scenario, initial nonrandom mating, single locus coding for the choosiness trait, few loci coding the ecological trait, high carrying capacity, high mutation rate) and we test the effects of weak (if *N*_male_ is intermediate) or strong (e.g., if *N*_male_ = 1) costs of choosiness on the evolution of choosiness. The high mutation rate implemented (*m_x_* = *m_c_* = 5.10^-2^) is offset by a small expected phenotypic variance at those traits (*s_x_* = *s_c_* = 0.002). This allows us to speed up the evolution of choosiness, while assessing a choosiness state close to its equilibrium value. Simulations end after 100,000 generations and 30 replicates are done for each combination of parameters.

### Selection gradient

To understand our simulation outputs, we then dissect the selective forces acting on choosiness by characterizing the direction of selection on choosiness in each deme. To do so, we numerically determine local selection gradients, defined by Geritz et al. (1997) as:

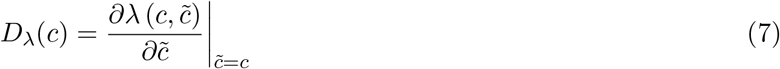

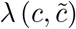 corresponds to the invasion fitness – i.e., the reproductive success – of a mutant with a choosiness trait *c* in a population with choosiness trait *c*. Therefore, *D*_λ_ describes how, in the vicinity of the resident mating strategy, a rare mutant’s strategy influences its (invasion) fitness. Choosiness (*c*) increases if *D*_λ_ is positive, and decreases if *D*_λ_ is negative. A convergence-stable intermediate equilibrium is achieved if *D*_λ_ = 0 and < 0.

To determine numerically *D*_λ_(*c*), we consider a population with choosiness trait *c* at its ecological equilibrium. we introduce 10% of mutants with a choosiness trait 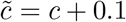, such that the alleles coding for 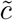 are in linkage disequilibrium with the ecological trait (this linkage typically arises after 3 generations and is assessed preliminary). After viability selection and reproduction, we measure the local frequencies of the allele coding for 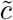 in the population and use them as a proxy of the invasion fitness 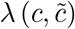. With analog simulations, we estimate λ (*c, c*). We then calculate the local selection gradient as:

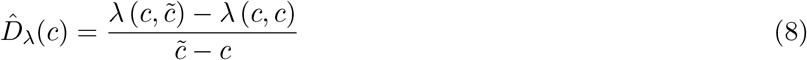

To characterize the direction of selection on choosiness in the entire population across all three demes, we estimate the global selection gradient as a weighted average of local selection gradients (calculated in the three demes). Following Wickman et al. (2017), the weights are the squares of the local population densities. Indeed, the intuitive approach of averaging selection across all members of the populations is wrong. Under weak selection, the resulting selection gradient takes the form of a weighted sum of selective effects, where the weights are the class frequencies and the reproductive values calculated in the resident population (Taylor and Frank, 1996). The reproductive value in a given class is the long-term contribution of individuals in this class to the future of the population relative to the contribution of other individuals in the population (Fisher, 1930). The reproductive value of individuals in a locality with high density is higher than in a locality with low density. Therefore, selection gradients at points in space where the population is abundant are disproportionately stronger than where it is rare. If the global selection gradient for choosiness is positive (resp. negative), stronger (resp. weaker) choosiness should evolve in the population. Additionally, we also decompose local selection gradients to infer how the frequency of the choosier allele changes as a result of viability selection (via viability costs) and sexual selection (via differential mating) in each deme.

Note that implementing a smaller initial fraction of mutant individuals and a smaller mutation size considerably increases the number of replicates (more than billions) and the computer runtime necessary to detect local selection in the main demes. Likewise, in simulations, different choosiness alleles can be present in the population at a given time step and the method computed here may look unrealistic in this regard. Nonetheless, the evolutionary dynamics of choosiness predicted by the global selection gradient predicts well the choosiness state reached (close to its equilibrium value) in our numerical simulations. We can then confidently use these statistics to dissect the selective forces acting on choosiness.

## RESULTS

### A weak cost of choosiness can favour the evolution of choosiness in parapatry

In the simulations, the population is characterized by two ecologically-specialized subpopulations, hereafter called ecotypes, distributed in parapatry. Over a large range of parameters *α*_T_ and *N*_male_ (relative size of the transition deme and maximum number of males encountered per female, respectively), populations evolve a positive choosiness trait – i.e., females mate preferentially with ecologically-similar males – (Fig. 2a), which favours the ecological differentiation between ecotypes (Fig. 2b). Therefore, strong nonrandom mating between ecotypes can evolve (Fig. 2c), reducing gene flow and leading to genetic differentiation at neutral loci (*F*_ST_ > 0, Fig. 2d) (see times series in Fig. S5). Since reproductive isolation reduces gene flow between ecotypes, the *F*_ST_ index reflects the strength of reproductive isolation between ecotypes. Importantly, choosiness is only intermediate; this result is in accordance with Servedio (2011) and Cotto and Servedio (2017), as explained in the section ’Selection on choosiness’ below (see also Appendix A).

**Figure 2:**
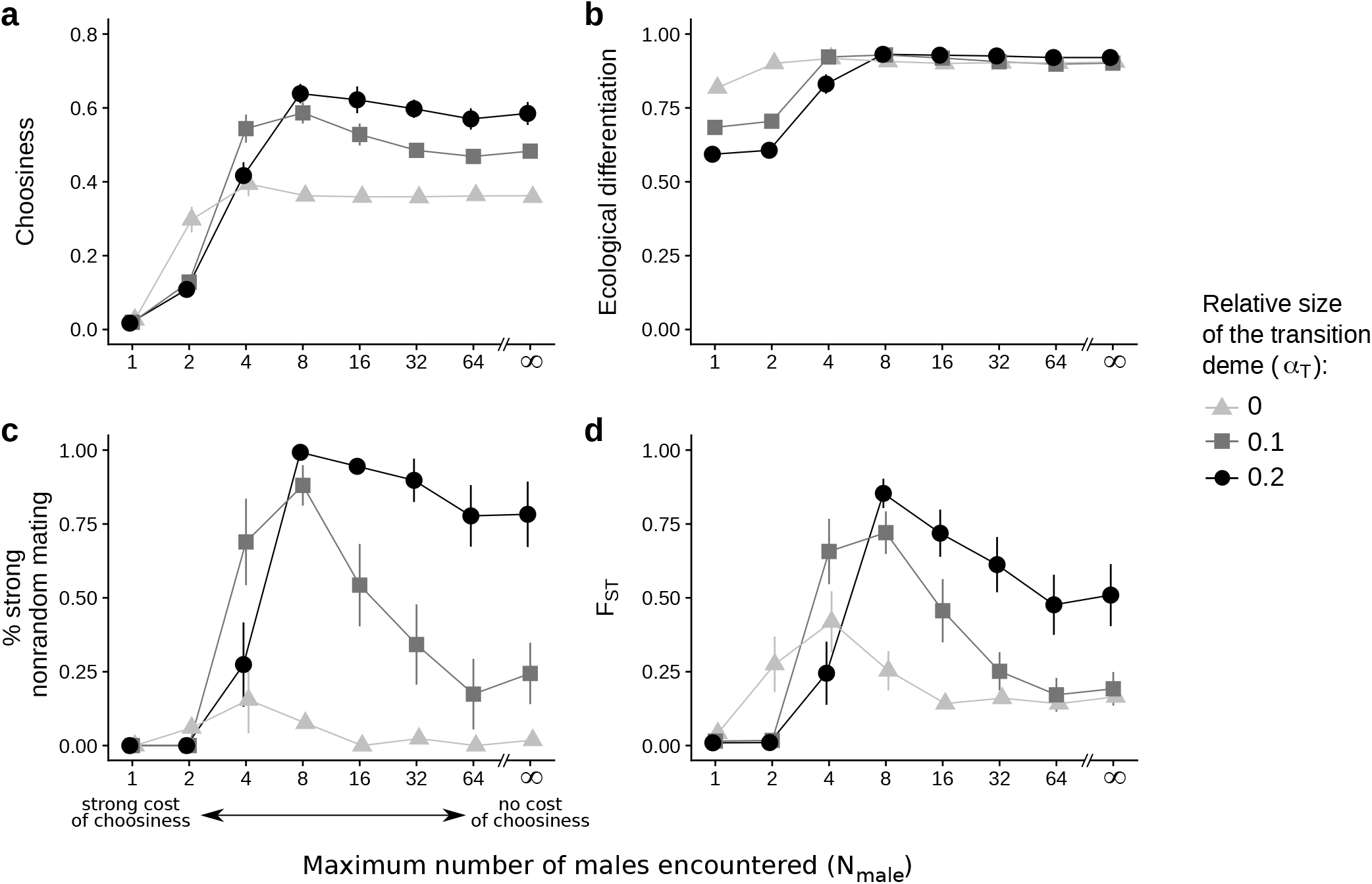
Ecological differentiation, nonrandom mating caused by choosiness and genetic differentiation with different geographical contexts and maximum numbers of males encountered per female. Statistics are measured from the traits and loci of individuals belonging to the two ecotypes (with ecological traits *x* ≤ 0.5 or *x* > 0.5). After 100, 000 generations, we record the choosiness in the population (a) and differences in ecological traits between ecotypes (b). To evaluate the overall strength of reproductive isolation, we measure both the percentage of runtime with strong nonrandom mating from generation 100, 000 to generation 110, 000 (c) and the genetic differentiation at neutral loci (*F*_ST_) (d). For each combination of parameters (*α*_T_, *N*_male_), the point and the error bars respectively represent the mean value of the statistic and its 95% confidence intervals. The x-axis is plotted on a logarithmic scale. *σ_s_* = 0.4.

In simulations with intermediate *N*_male_ values, choosy females face a small risk to remain unmated, but despite this weak cost of choosiness, assortative mating evolves and reduces gene flow between ecotypes (*F*_ST_ > 0; Fig. 2). Interestingly, with such weak cost of choosiness, stronger choosiness evolve compared to simulations with no cost of choosiness (*N*_male_ = ∞; when all females eventually find a mate) (Fig. 2a). While this increase in choosiness may appear anecdotal at first glance, it actually leads to stronger assortative mating (Fig. 2c) and much higher genetic differentiation (Fig. 2d) (see also times series in Fig. S5).

In simulations with very low *N*_male_ values, all choosy females have very high chance to remain unmated. With such strong cost of choosiness (e.g., *N*_male_ = 2), evolution of choosiness is severely restricted, and with a maximal cost of choosiness (*N*_male_ = 1), no choosiness evolves and gene flow is unrestricted (*F*_ST_ = 0; Fig. 2).

Costs of choosiness affect the evolution of choosiness depending on the strength of disruptive viability selection (inversely proportional to *σ_s_*). If viability selection is very strong (resp. very weak), choosiness is strongly (resp. weakly) favoured, so a weak cost of choosiness has no effect on the evolution of choosiness (for *σ_s_* ≤ 0.2 and *σ_s_* ≥ 0.5, Fig. S6). The effects of costs also depends on the relative size of the transition deme (*α*_T_). Under weak viability selection, a weak cost of choosiness strongly favours the evolution of choosiness if the transition deme is large (for *α*_T_ > 0 if *σ_s_* ≥ 0.4, Fig. S6). Under strong viability selection, a cost of choosiness strongly favours the evolution of choosiness if the transition deme is small (for *α*_T_ = 0 if *σ_s_* ≤ 0.3, Fig. S6). If choosiness is costly, strong choosiness may evolve if the transition deme is small and not if the transition deme is large (e.g., for *N*_male_ = 2 in Figs. 2 and S6). Note that we get similar results if we vary the size of the transition deme, without changing the size of the main demes (Fig. S7).

### Selection on choosiness

We aim at understanding how the cost of choosiness affects selection on choosiness. We here dissect in each deme the different selective forces (viability and sexual selection) acting on choosier alleles (for *α*_T_ = 0.1, Fig. 3). Note that we can draw the same conclusions from the selection gradients obtained in ecological settings with a different relative size of the transition deme (*α*_T_ = 0 or 0.2; see Figs. S8 and S9).

**Figure 3:**
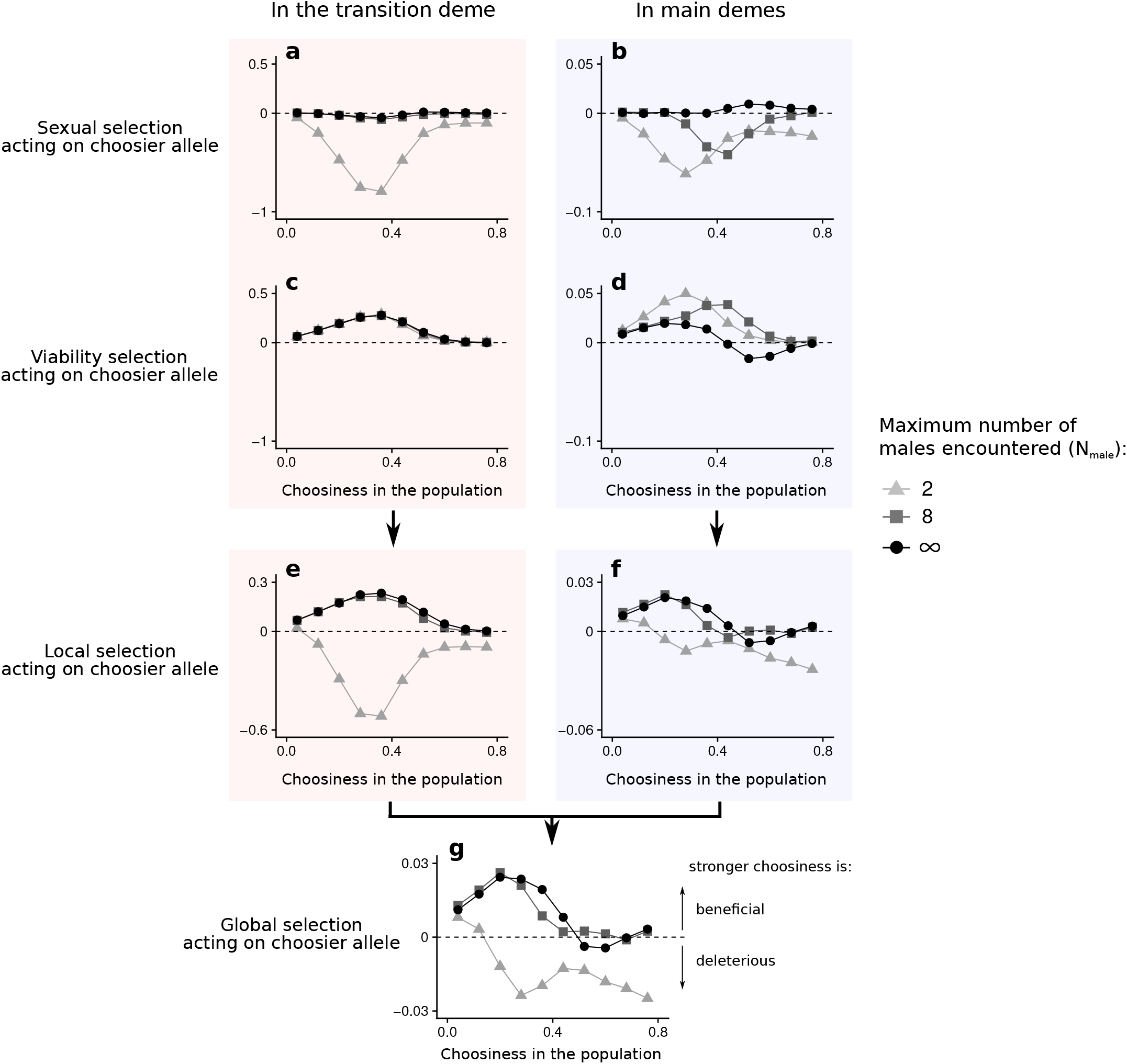
Local selective forces acting on choosiness in the transition deme (a, c, e) and in main demes (b, d, f), and global selection acting on choosiness (g). After the introduction of 10% of mutants with a choosier allele than the one carried by all other individuals in the population, local selection gradients are measured as changes in frequencies of these choosier alleles over one generation (e-f). Sexual selection (resp. viability selection) acting on alleles coding for stronger choosiness is measured as their change in frequencies during the reproduction phase (resp. viability selection phase) (a-d). Following Wickman et al. (2017), we estimate the global selection gradient as a weighted average of local selection gradients (g). Note that scales of the y-axis in the left and right column are different. *σ*_s_ = 0.4, *α*_T_ = 0.1.

#### *With no cost of choosiness* (N_*male*_ = ∞)

We first consider the situation where all females eventually find a mate (black points in Figs. 3 and 4). In the transition deme, alleles coding for strong choosiness are associated with locally favoured alleles at the ecological trait loci (i.e., with alleles coding for extreme ecological phenotypes x close to 0 or 1) (linkage disequilibrium caused by selection, Fig. 4a,c). Therefore, selection on the ecological trait indirectly acts on choosiness via this linkage disequilibrium. Positive frequency-dependent sexual selection – i.e., high mating success of males whose ecological phenotype is in high frequency locally – weakly inhibits or favours the evolution of choosiness in the transition deme, depending on the relative proportions of ecological phenotypes (Fig. 3a). More importantly, disruptive viability selection – i.e., low survival of males with intermediate ecological phenotypes – strongly favours the evolution of choosiness in the transition deme (Fig. 3c). In the transition deme, selection therefore favours the evolution of strong choosiness (Fig. 3e).

**Figure 4:**
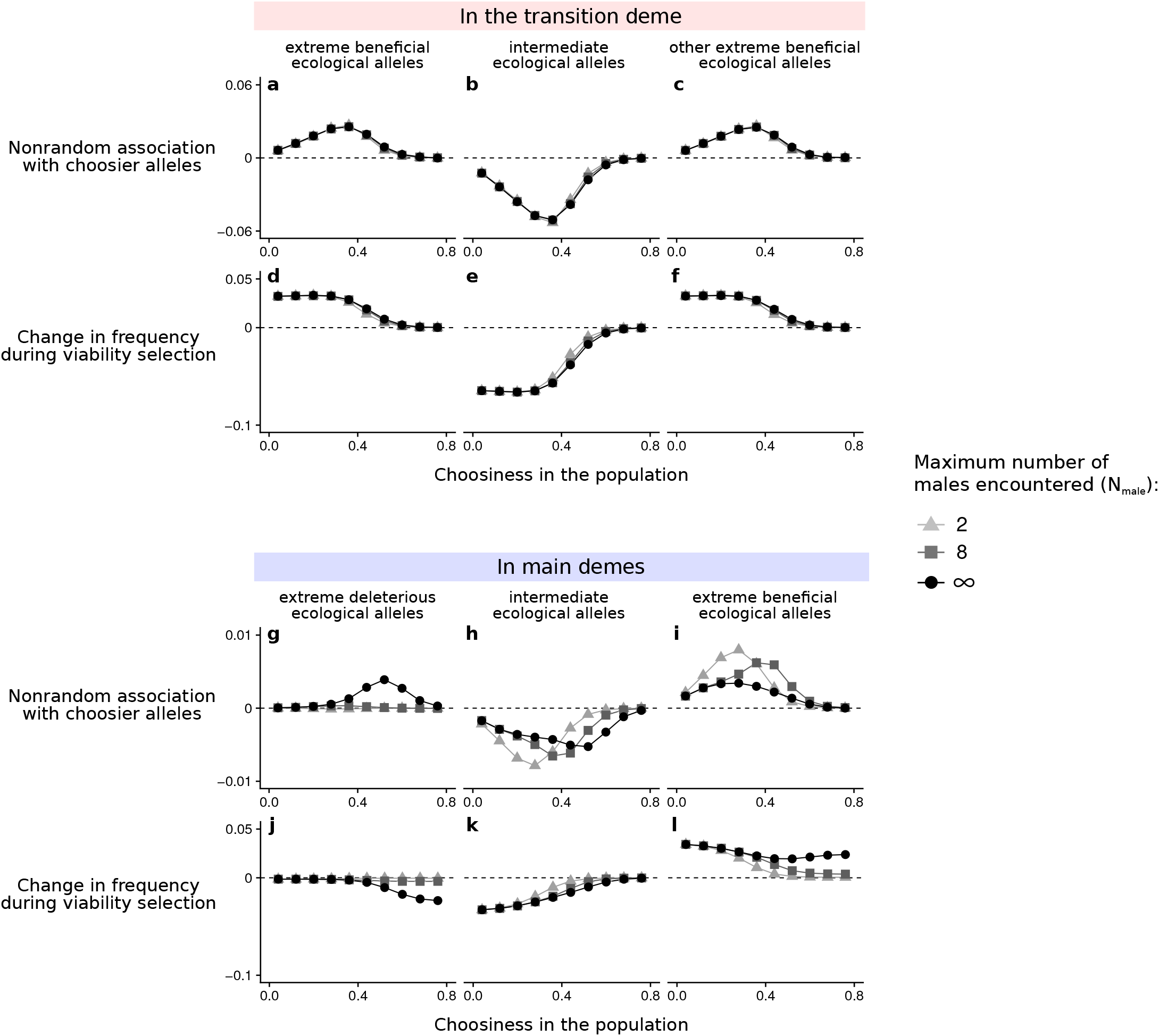
Linkage disequilibrium between choosiness and ecological loci after reproduction and change in frequency of ecological categories via direct viability selection in the transition deme (a-f) or in main demes (g-l). Individuals are categorized according to their ecological alleles (coding for: *x* < 0.33, 0.33 ≤ *x* ≤ 0.66 or *x* > 0.66). Three generations after implementing mutant alleles at the choosiness locus, linkage disequilibrium between choosiness and ecological loci is measured as the nonrandom association of alleles coding for stronger choosiness with each ecological category after reproduction. We also measure the change in frequency of each ecological category during viability selection. Altogether, these graphs allow us to infer how viability selection indirectly acts on choosiness via linkage disequilibrium. In particular, for *N*_male_ = ∞, indirect viability selection inhibits the evolution of choosiness in main demes via a positive linkage between alleles coding for stronger choosiness and locally unfavoured alleles at the ecological locus. This positive linkage vanishes for *N*_male_ = ∞, and so does indirect viability selection inhibiting strong choosiness. *σ_s_* = 0.4, *α*_T_ = 0.1.

In the main demes, alleles coding for strong choosiness are associated with locally favoured alleles at the ecological trait loci (here, with alleles coding for the extreme ecological phenotype that is locally favoured) (Fig. 4i). Just like in the transition deme, positive frequency-dependent sexual selection weakly inhibits or favours the evolution of choosiness, depending on the distribution of ecological phenotypes (Fig. 3b). More importantly, alleles coding for strong choosiness are strongly associated not only with locally *favoured* alleles but also with locally *unfavoured* alleles at the ecological trait loci (i.e., leading to a negative association with alleles coding for an intermediate ecological trait, Fig. 4g-i; as shown with a simple two-deme population genetic model in Appendix A). Indeed, females with a rare ecological phenotype mostly mate assortatively if they are very choosy (whereas they mostly mate with locally adapted males, because they are common, if they are only weakly choosy). This linkage leads to the removal of alleles coding for strong choosiness via indirect viability selection (Fig. 4j). In other words, viability selection indirectly inhibits the evolution of strong choosiness (Fig. 3d). In main demes, selection therefore favours the evolution of intermediate choosiness (Fig. 3f).

In the absence of cost of choosiness, strong choosiness is favoured in the transition deme (Fig. 3e) but not in the main demes (via indirect viability selection; Fig. 3f). Overall, the global selection gradient predicts that for *α* = 0.1, only intermediate choosiness can evolve (where the black line crosses the horizontal dashed line in Fig. 3g). This explains why in our numerical simulations, intermediate choosiness evolves if the transition deme is small, and strong choosiness evolves if the transition deme is large (Fig. 2). These results without a cost of choosiness are consistent with the results of Cotto and Servedio (2017) (see also Appendix A).

#### *With a cost of choosiness* (N_*male*_ ≠ ∞)

A cost of choosiness (when choosy females risk to remain unmated) induces sexual selection inhibiting choosiness via differential *female* mating success (Fig. 3a-b). This additional sexual selection pressure is therefore distinct from positive frequency-dependent sexual selection caused by differential *male* mating success.

In the transition deme, a strong cost of choosiness induces very strong sexual selection inhibiting the evolution of choosiness (e.g., for *N*_male_ = 2), whereas a weak cost of choosiness induces weak sexual selection compared to the strength of viability selection (e.g., for *N*_male_ = 8) (Fig. 3a). Additionally, indirect viability selection strongly favours the evolution of choosiness in the transition deme (Fig. 3c). Therefore, only a strong cost of choosiness can inhibit the evolution of choosiness in the transition deme (Fig. 3e).

In the main demes, a cost of choosiness induces sexual selection inhibiting the evolution of choosiness (Fig. 3b), but even with a strong cost (*N*_male_ = 2), this sexual selection pressure is not particularly strong, because locally adapted females often find a matching mate at the first encounter (Fig. 3b). Crucially, a cost of choosiness also modifies the local population assembly on which viability selection acts. In particular, choosy females with a rare ecological phenotype have little chance to find preferred (matching) males and often remain unmated. Consequently, a cost of choosiness weakens the association between alleles coding for strong choosiness and locally *unfavoured* alleles (in low frequency) at the ecological trait loci (Fig. 4g). A cost of choosiness can therefore cause a positive linkage disequilibrium between alleles coding for strong choosiness and locally *favoured* alleles at the ecological trait loci (Fig. 4i), and this effect leads to indirect viability selection favouring strong choosiness in main demes (Figs. 3d and 4j-l). Notably, in Appendix A, we show that a cost of choosiness can cause such positive linkage disequilibrium in a simple two-deme population genetic model. Therefore, in main demes, a cost of choosiness induces additional direct sexual selection inhibiting choosiness, and causes indirect viability selection favouring choosiness. Overall, in the main demes, a strong cost of choosiness inhibits the evolution of choosiness (for *N*_male_ = 2, Fig. 3f). A weak cost of choosiness, however, inhibits the evolution of intermediate choosiness (via additional sexual selection) but favours the evolution of strong choosiness (via indirect viability selection) (for *N*_male_ = 8, Fig. 3f; the dark grey curve crosses the x-axis twice).

With a strong cost of choosiness, choosiness is deleterious in the transition deme and in the main demes (for *N*_male_ = 2, Fig. 3e-f). With a weak cost of choosiness, however, choosiness is favoured in the transition deme (via indirect viability selection which overcome sexual selection caused by the cost of choosiness across the selection gradient) and indirect viability selection now favours the evolution of strong choosiness in the main demes (for *N*_male_ = 8, Fig. 3e-f). How can intermediate choosiness and then strong choosiness evolve from a state of random mating when there is a weak cost of choosiness? While the evolution of intermediate choosiness is often hindered in the main demes due to the cost, intermediate choosiness can initially evolve because it is strongly favoured in the transition deme. Then, strong choosiness is favoured not only in the transition deme, but also in the main demes because indirect viability selection favours the evolution of strong choosiness when there is a weak cost choosiness. This explains why a weak cost of choosiness can favour the evolution of strong choosiness in our simulations (Fig. 2).

Moreover, unlike in main demes, a strong cost of choosiness induces strong sexual selection against choosiness in the transition deme (for *N*_male_ = 2, Fig. 3a). This explains why, if choosiness is very costly, strong choosiness can evolve if the transition deme is small and not if the transition deme is large (Fig. 2).

## DISCUSSION

In a parapatric context of speciation, the evolution of assortative mate choice is driven by non-intuitive selective forces. We have shown that in parapatry, costs of choosiness have a non-linear effect on the evolution of choosiness, and therefore on reproductive isolation. Unsurprisingly, a strong cost of choosiness taking the form of a high risk of remaining unmated inhibits the evolution of choosiness in parapatry. By constrast, a moderate cost of choosiness, which is mainly incurred by immigrants, can *favour* the evolution of choosiness. This counterintuitive prediction relies on the effects of this cost on local sexual and viability selection in different localities across space. Based on the predictions of our model, the width of the ecological transition zone, the strength of fitness costs, and the ecology of mating all prove to be important when examining the likelihood of parapatric speciation.

Previous models of parapatric speciation have not accounted for the possibility that females may suffer from delayed mating if they are are too choosy (Kawata et al., 2007; Leimar et al., 2008; Ispolatov and Doebeli, 2009; Servedio, 2011; Cotto and Servedio, 2017), while this ‘cost of choosiness’ should be high for female immigrants in foreign environments (as argued by Servedio and Hermisson, 2020). In our study, we model the deleterious effect of delayed mating as a risk of remaining unmated (a situation observed in nature; Reynolds and Gross, 1990; Milinski et al., 1992; Heubel et al., 2008; Scott et al., 2020). We show that in parapatry this cost of choosiness induces sexual selection which inhibits the evolution of choosiness just like in sympatry (Bolnick, 2004; Schneider and Bürger, 2006; Bürger et al., 2006; Kopp and Hermisson, 2008). Interestingly, however, this cost of choosiness also changes linkage disequilibria, and thereby leads to indirect viability selection favouring the evolution of strong choosiness in parapatry.

In parapatry, selection favours intermediate choosiness because strong choosiness creates a positive genetic association between choosiness and locally *disfavoured* ecological alleles (Servedio, 2011; Cotto and Servedio, 2017). In our study, we showed that a cost of choosiness taking the form of a risk of remaining mating can weaken this genetic association, thereby indirectly favouring the evolution of strong choosiness. This is because outside of the ecological transition zone, ecotype frequencies are unbalanced, so the difficulty of finding a matching mate mostly affects female immigrants, which also suffer viability costs due to maladaptation. Therefore, maladapted females tend not to mate at all, preventing the build-up of linkage disequilibrium between maladapted ecological alleles and strong choosiness. As a result, viability selection no longer removes alleles coding for strong choosiness via linkage disequilibrium. By lessening the negative effect of indirect viability selection on choosiness, a cost of choosiness can therefore favour the evolution of strong choosiness through indirect viability selection. In other words, outside of the ecological transition zone, sexual selection changes the phenotype/allele frequencies on which viability selection acts (as shown without a cost of choosiness by Cotto and Servedio, 2017); this leads to this counterintuitive effect of the costs of choosiness on the evolution of choosiness.

Local selection on choosiness varies spatially as ecotypes can coexist in some localities (as in the ecological transition zone) or exclude themselves in others. A weak cost of choosiness (for *N*_male_ = 8 in our simulations) does not change net selection on choosiness in the ecological transition zone, because sexual selection induced by this cost is always offset by strong indirect viability selection favouring choosiness. On the contrary, a weak cost of choosiness affects net selection acting on choosiness outside the ecological transition zone. In particular, sexual selection induced by such cost also generates indirect viability selection favouring strong choosiness. Consequently, the overall effect of a weak cost of choosiness may change according to the relative importance of the transition zone and the main demes, where selection on choosiness may act in opposite directions depending on the strength of viability selection. In other words, the abruptness of the spatial ecological transition – i.e., the extent of range overlap between divergent ecotypes – and the fitness effects of ecological adaptation determine the evolution of choosiness when the cost of choosiness is weak.

A more detailed look into alternative genetic architectures (e.g., “two-allele mechanisms”, physical linkage among choosiness loci) could conceivably change the effects of costs of choosiness on reproductive isolation in parapatry, and should therefore be investigated in future theoretical studies. In a two-deme population genetic model accounting for a two-allele mechanism, Servedio and Burger (2014) found that a cost of choosiness can in some cases lead to stronger differentiation, but whether stronger choosiness can evolve remains to be formally investigated.

Without a cost of choosiness, range overlap between ecotypes has been shown to affect the evolution of assortative mating in parapatry. In our model, wide range overlap between ecotypes leads to the loss of polymorphism at the ecological trait because of positive frequency-dependent sexual selection (Kirkpatrick and Nuismer, 2004; Otto et al., 2008; Cotto and Servedio, 2017) (see Fig. S4), while narrow range overlap gives scope for indirect viability selection inhibiting strong choosiness (Servedio, 2011; Cotto and Servedio, 2017). Without a cost of choosiness, reproductive isolation between divergent ecotypes is therefore the strongest for intermediate ecological transitions (as previously shown by Doebeli and Dieckmann, 2003; Cotto and Servedio, 2017). Here we show that this result changes qualitatively when choosiness is costly. Since costs of choosiness have a stronger effect in the ecological transition zone (where the ranges of the two ecotypes overlap) than outside of the ecological transition zone (where they do not), strong costs might impede the evolution of strong choosiness for a wide range overlap but not for a narrow range overlap. Assessing the strength of the cost of choosiness (if any) is therefore of prime importance to predict how range overlap would affect the evolution of assortative mating.

Hybrid zones are transition zones between the geographical ranges of closely related taxa where hybrids are produced (Hewitt, 1988). Hybrid zones are often represented as narrow strips separating much larger regions in which the two taxa reside, like in our model. When the gradient is environmental (adaptation to resources which are unevenly distributed for instance), the width of a hybrid zone depends on the change in environmental conditions. However, other hybrid zones are ‘tension zones’ maintained by a balance between dispersal and selection against hybrids (Slatkin, 1973; Mallet and Barton, 1989), and the width of such tension zones therefore results from this balance. Mallet et al. (1990) have used the width of multiple tension zones to estimate gene flow between races of *Heliconius erato* and those of *Heliconius melpomene.* Yet, in this study, gene flow was approximated by dispersal estimates, whereas actual gene flow also depends on pre- and post-zygotic reproductive barriers. We hope to stimulate similar empirical investigations on multiple hybrid zones, but this time precisely testing such a link between hybrid zone width (or the steepness of the environmental gradient) and actual gene flow (that could be assessed from the F_ST_). To our knowledge, only Seehausen et al. (2008) empirically showed such link between gradient steepness and gene flow (approximated by differentiation). Specifically, they showed that differentiation in cichlid fishes does not occur on very steep environmental light gradients. In that case, our predictions suggest that cost of choosiness may be absent or weak. Otherwise, differentiation would have been favoured on very steep gradient. In the light of our theoretical predictions, similar empirical investigations would considerably strengthen our understanding of the role of the spatial dimension of ecological changes in parapatric speciation.

We acknowledge that migration is usually more complex than a simple ‘reciprocity rule’ as implemented in our model. In normal “tension zone” dynamic, there is often less migration out of the hybrid zone than into it. In such context, selection within the zone is likely to have a different impact than in the case of reciprocal migration. Likewise, our representation of space is still very simplified, since distributions of ecotypes along ecological gradients can form a diversity of patterns, from continuous gradations from one ecotype to another to patchy distributions of ecotypes called mosaic hybrid zones (Harrison and Larson, 2016). In a context of a steep ecological gradient, our results should not depend on the ecotype distribution inside the transition region at a smaller scale. On the contrary, if the transition region is large, our predictions may not be robust to a context of mosaic hybrid zone. In that case, the relative proportion of the two taxa is always skewed in favour of one or the other taxon, even in the transition region. Therefore, local selection acting in each ‘patch’ of the mosaic may be similar to the one acting at the tails of the gradient where ecotype frequencies are unbalanced. The predictions of our model suggest that, under a context of mosaic hybrid zone, assortative mating should readily evolve if choosiness is costly. It may explain why reproductive isolation between *Timema cristinae* (stick insect) ecotypes – distributed in such mosaic hybrid zone – is incomplete in that complete speciation has not occured (assuming that the cost of choosiness is weak or absent) (Nosil, 2007).

In a recent theoretical study, Irwin (2020) explicitly considered a diverging population distributed along a ‘tension zone’ with all individuals having the same level of choosiness. He showed that costs of choosiness can prevent blending between incipient species, and can maintain a parapatric distribution (see also M’Gonigle et al., 2012). This adds support to the idea highlighted in our study that costs of choosiness can promote parapatric speciation, this time by maintaining the population distribution along an ecological gradient. We did not consider the feedback of the evolution of choosiness on the width of the hybrid zone; whether this effect is strong enough as choosiness evolves warrants further theoretical investigation.

In the light of the predictions of our model, the maximum number of males that females can encounter in their lifetime is a key factor for the evolution of reproductive isolation in parapatry. Females usually cannot sample all males within a population, because of such factors as environmental constraints on the timing of reproduction and predation risks associated with searching (Morris, 1989; Wagner, 1998). However, it is difficult to estimate how many males females can evaluate in nature. The number of males evaluated by females prior to breeding has long been the best estimate available (Bolnick and Fitzpatrick, 2007). However, movement pattern has also been acknowledged to determine the potential for female mate choice to drive sexual selection (Dunn and Whittingham, 2007; Duval and Kapoor, 2015). Therefore, empirical data on how free-living females move among and select mates may provide information on potential costs of choosiness. For instance, Kamath and Losos (2018) managed to estimate encounters between potential mates from mark-recapture data of male and female *Anolis sagrei* lizards. Applying this method along ecological gradients on different ecotypes may provide good estimates of potential costs of choosiness. Indeed, from the estimation of encounter estimates within vs. between ecotypes, the risk of remaining unmated associated to choosiness may easily be assessed. In our model, females do not adjust their level of choosiness in response to the abundance of males from their own type (as shown by Willis et al., 2011). In this case, choosy immigrant females would keep mating with local common males and evolution of strong choosiness would not inhibit hybridization between ecotypes (as noted by Kopp and Hermisson, 2008). Such choosiness subject to adjustement may be best considered partial rather than strong.

Overall, our theoretical model sheds new light on the role of spatial dimension of ecological change in parapatric speciation. We show that, in parapatry, sexual selection pressures can have very counterintuitive effects on trait evolution by locally affecting the genotypic frequencies on which viability selection acts. In parapatry, a cost of choosiness (when choosy females may risk to remain unmated) can even favour the evolution of choosiness and subsequent reproductive isolation. Consequently, the effect of spatial dimension of ecological change on speciation also depends on the strength of costs of choosiness. Our study should therefore stimulate further empirical research on the link between spatial structure, costs of choosiness and reproductive isolation.

## ACKNOWLEDGMENTS

We thank M. Kopp, C. Thébaud, E. Porcher, C. Smadja, M. Servedio, B. Lerch, and K. Xu for insightful discussions and for comments on the manuscript. Simulations were performed on the computing cluster of the Montpellier Bioinformatics Biodiversity (MBB) platform. This research was supported by grants from the Swiss National Science Foundation and from the National Science Foundation (to T.G.A.).

## DATA AVAILABILITY STATEMENT

The code for the simulations will be deposited at Dryad Digital Repository.

