## Supplementary Figures for "Costs of choosiness can promote reproductive isolation in parapatry"

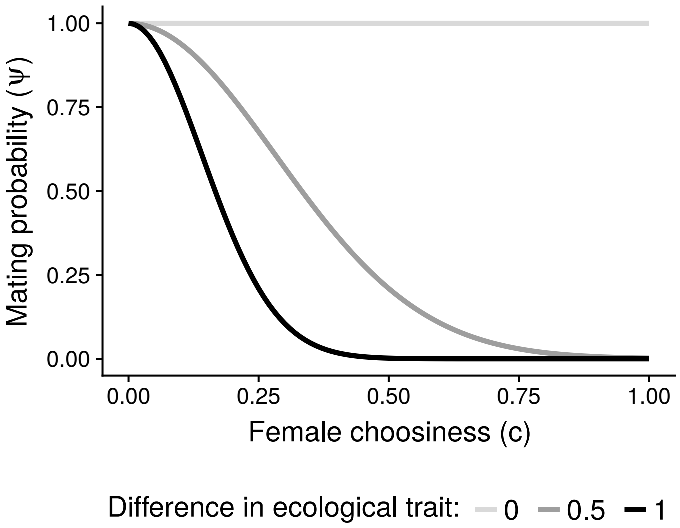


Figure S1: Mating probability between a female with ecological trait $x_f$ and choosiness $c$ and a male with ecological trait $x_m$. The difference in ecological trait between the male and the female $\mid x_f - x_m \mid$ is 0, 0.5 or 1. When a female is choosy (high choosiness trait $c$), she mates with males with a similar ecological trait ($\Psi = 1$ if $\mid x_f - x_m \mid = 0$), and she rejects ecologically-different males ($\Psi < 1$ if $\mid x_f - x_m \mid = 1$).

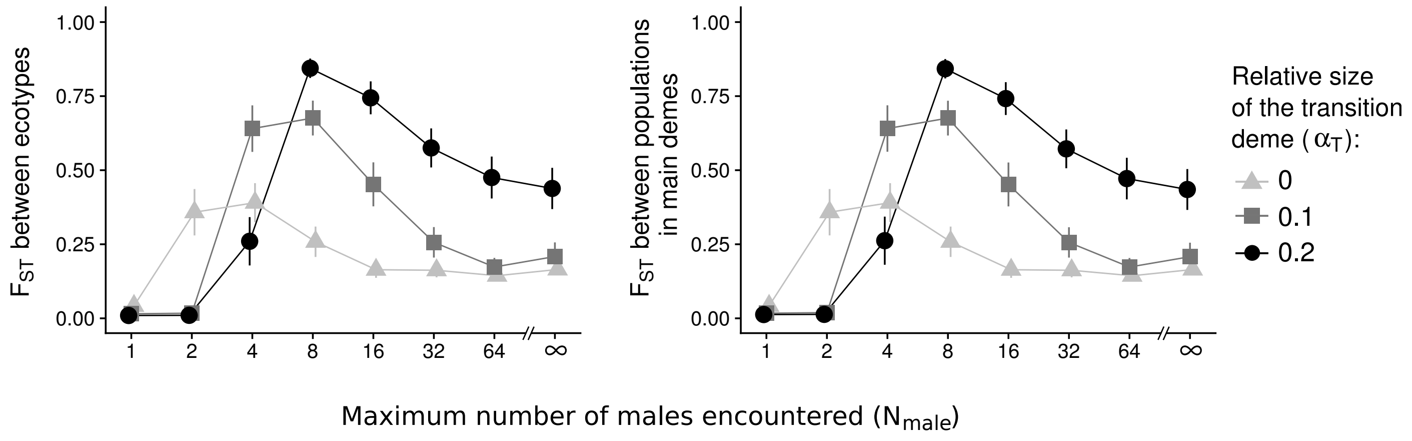


Figure S2: Genetic differentiation between ecotypes or between population from the two main demes. See Fig. 2 for details. Whether the $F_{\mathrm{ST}}$ statistics is calculated from the two ecotypes (with either $x \leq 0.5$ or $x > 0.5$) or the populations from the two main demes does not matter. $\sigma_{\mathrm{s}} = 0.4$.

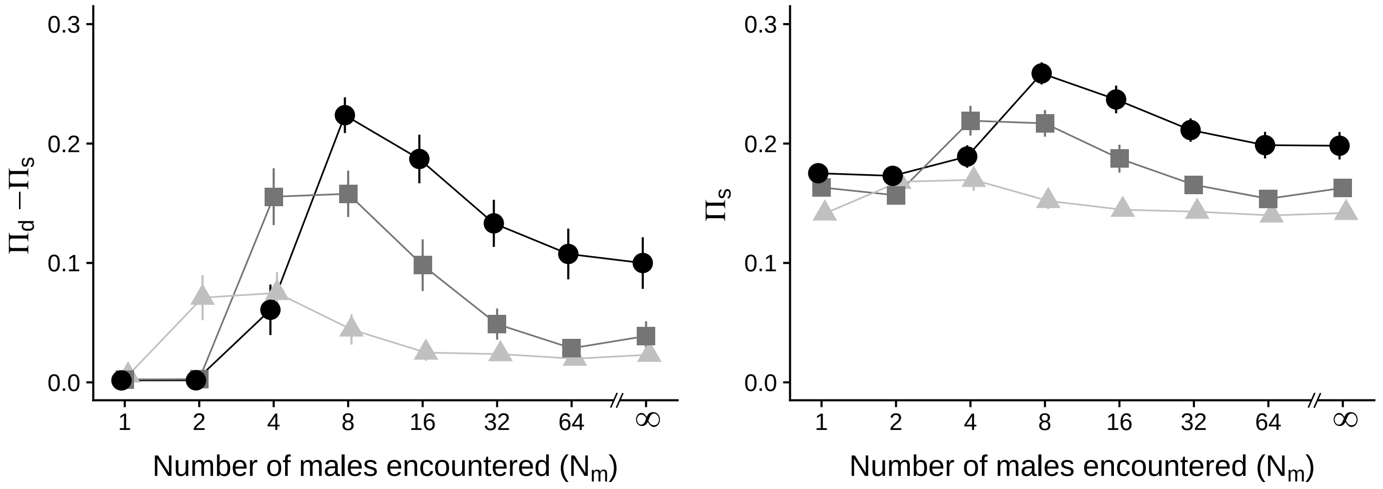


Figure S3: Numerator and denumerator of the genetic differentiation measure, $F_{\mathrm{ST}} = \frac{\Pi_d - \Pi_s}{\Pi_s}$. See Fig. 2 for details. We get qualitatively the same result when accounting for absolute differentiation, $\Pi_d - \Pi_s$. $\sigma_{\mathrm{s}} = 0.4$.

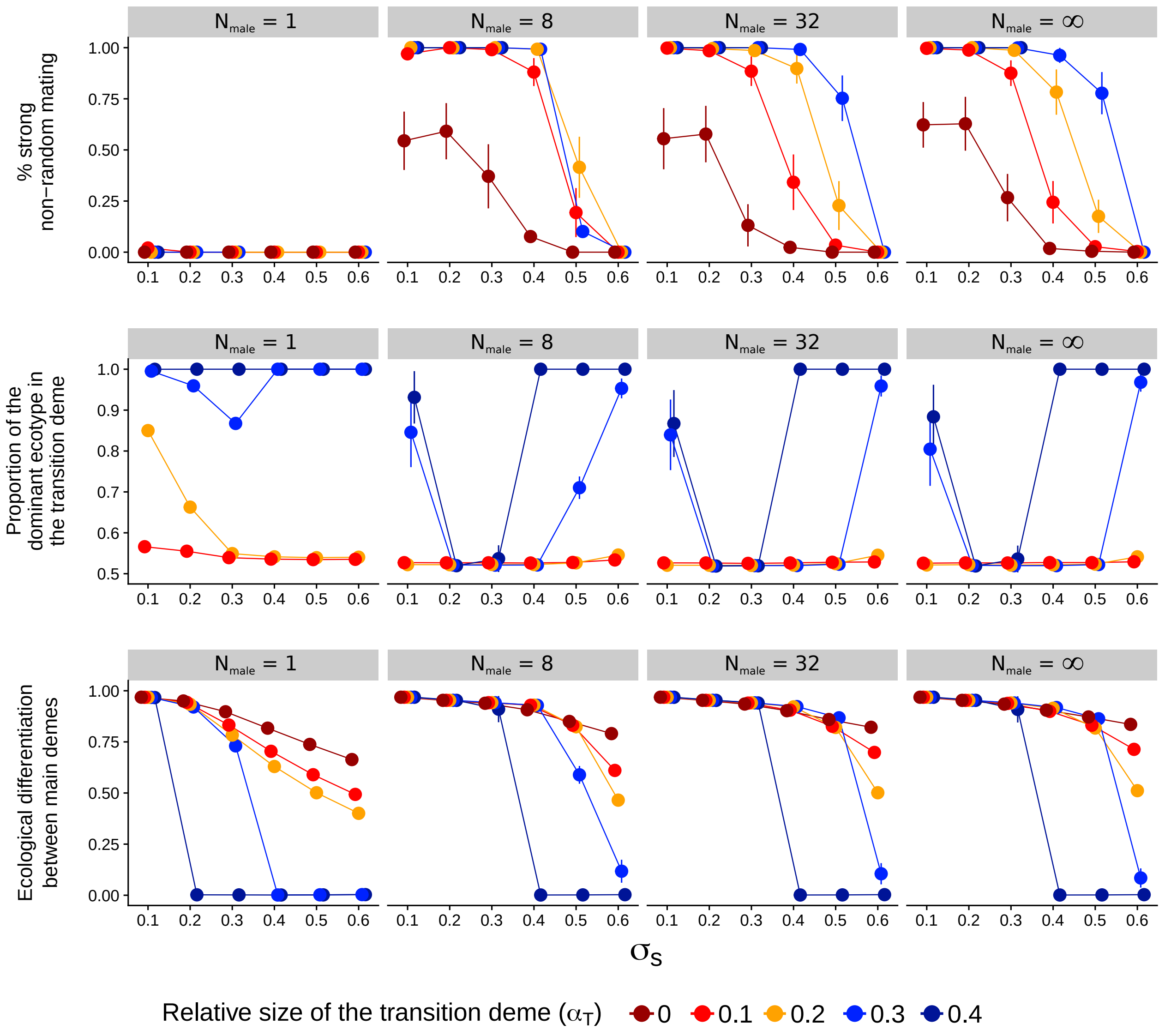


Figure S4: Nonrandom mating and ecological composition in the transition deme and in main demes, with different geographical contexts and strengths of viability selection (inversely proportional to $\sigma_s$). After 100,000 generations, percentage of strong nonrandom mating is recorded to evaluate the strength of reproductive isolation between ecotypes (with either $x \leq 0.5$ or $x > 0.5$). The dominant ecotype correspond to the more frequent ecotype in the transition deme. If the proportion of the dominant ecological type is close to 0.5 (resp. 1), the population in the transition deme is polymorphic (resp. monomorphic). The ecological differentiation between main demes is measured by comparing the ecological traits of individuals belonging to the different main demes. Polymorphism at the ecological locus is lost if there is no ecological differentiation between the main demes. For each combination of parameters $(\alpha_T, \sigma_s)$, the point and the error bars respectively represent the mean value of the statistic and its 95% confidence intervals. If the transition deme is too large ($\alpha_T \geq 0.3$), polymorphism can be lost or the transition deme can be polymorphic. If disruptive viability selection is too weak ($\sigma_s \geq 0.6$), assortative mating does not evolve, whereas, if disruptive viability selection is too strong ($\sigma_s \leq 0.1$), the transition deme is monomorphic for $N_{\text{male}} = 1$. To analyze the forces acting on the evolution of choosiness in populations with similar spatial structures (with a polymorphic transition deme), we therefore choose to restrict our main analyses to the ranges of parameters $\sigma_s \in [0.2, 0.5]$ and $\alpha_T \in [0, 0.2]$.

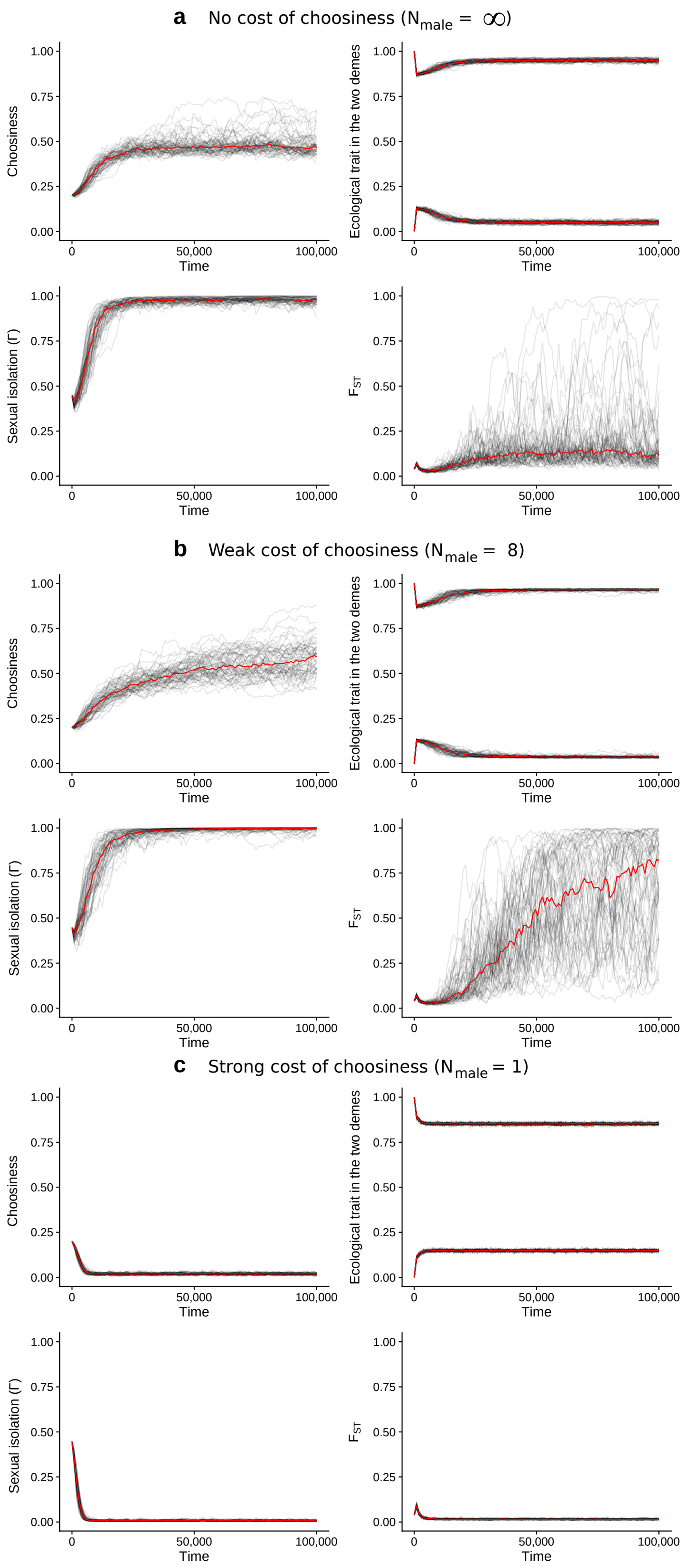


Figure S5: Time series. Each line correspond to the time series for one of the 100 replicates. The red line shows the median value of each statistics at each time step considered. With a weak cost of choosiness, stronger choosiness can evolve than without a cost of choosiness, leading to higher reproductive isolation and higher genetic differentiation. $\sigma_{\mathrm{s}} = 0.4$, $\alpha_{\mathrm{T}} = 0.1$.

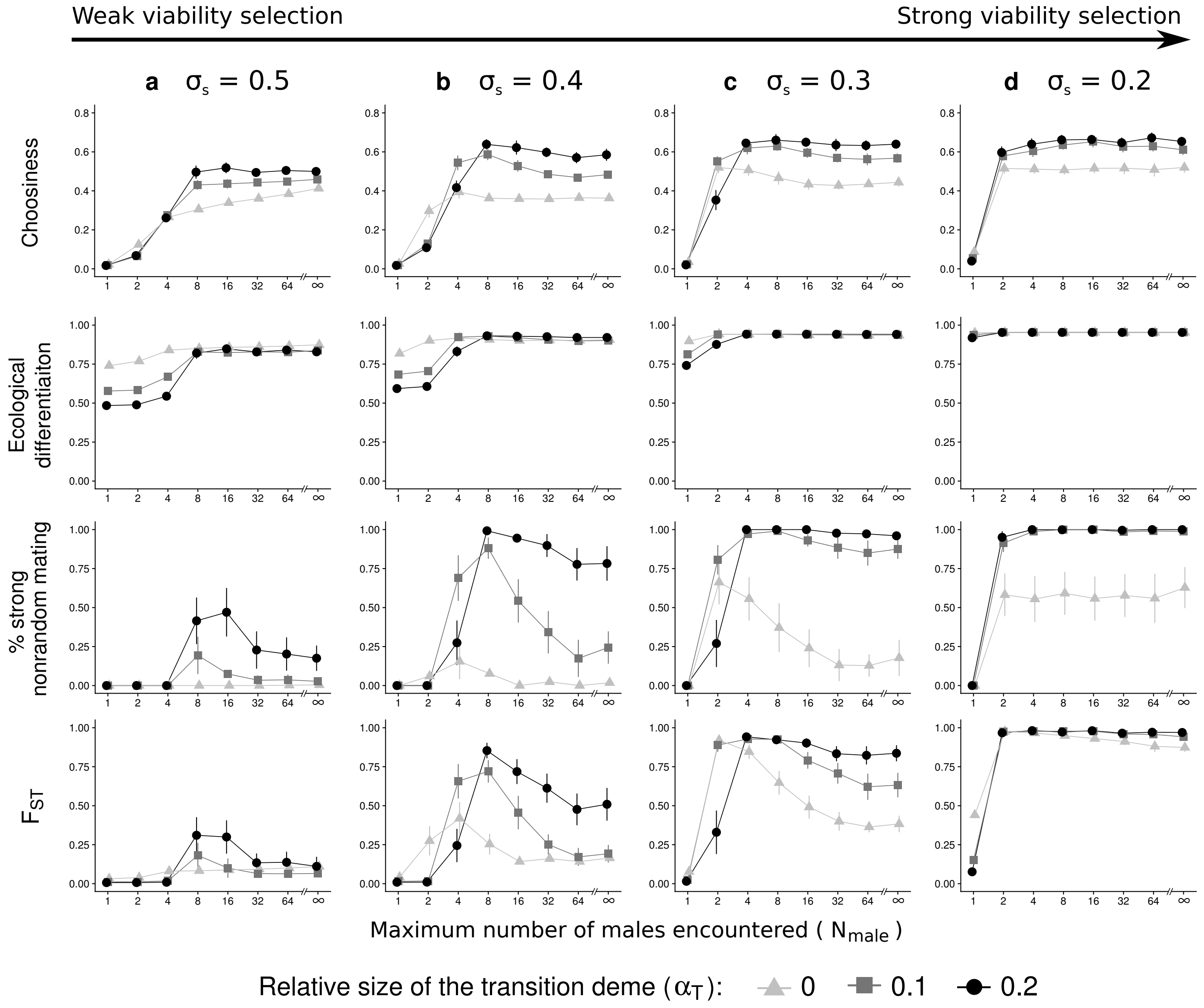


Figure S6: Effect of the strength of viability selection (inversely proportional to $\sigma_\mathrm{s}$) on the evolution of choosiness and subsequent reproductive isolation between ecotypes. See Figure 2 for details. Strong disruptive viability selection favours the evolution of choosiness, which leads to strong nonrandom mating, and favours genetic differentiation at neutral loci. Contrary to a strong cost of choosiness ($N_\mathrm{male} = 1$), a weak cost of choosiness (intermediate $N_\mathrm{male}$) favour the evolution of nonrandom mating and genetic differentiation among ecotypes, especially if disruptive viability selection is intermediate ($\sigma_\mathrm{s} = 0.3$ and 0.4). Additionally, with a weak cost of choosiness, uncommon females that are partially choosy have little chance to mate, reducing hybridization rate between ecotypes. Therefore, under some combinations of parameters, $F_\mathrm{ST}$ index is increased for intermediate $N_\mathrm{male}$ without evolution of stronger choosiness (e.g., for $\sigma_\mathrm{s} = 0.2$ and $\alpha_\mathrm{T} = 0$).

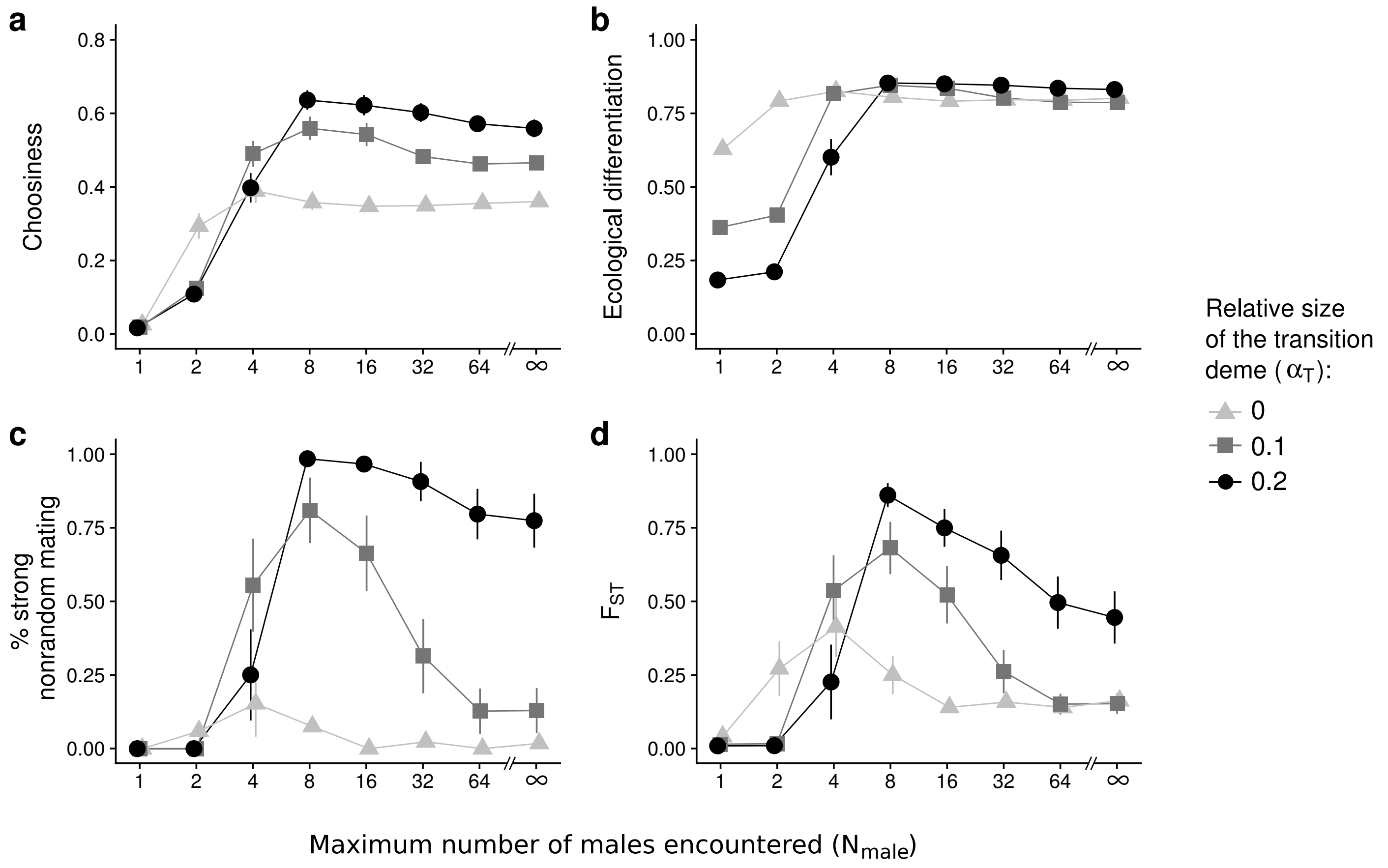


Figure S7: Ecological differentiation, nonrandom mating caused by choosiness and genetic differentiation with different geographical contexts **(with constant $\mathbf{K_1 = K_2 = 5,000}$)** and maximum numbers of males encountered per female. Statistics are measured from the traits and loci of individuals belonging to the two ecotypes (with ecological traits $x \leq 0.5$ or $x > 0.5$). After 100,000 generations, choosiness in the population (a) and differences in ecological traits between ecotypes (b) are recorded. To evaluate the strength of reproductive isolation, both the percentage of strong nonrandom mating (c) and the genetic differentiation at neutral loci ($F_{\mathrm{ST}}$) (d) are measured. For each combination of parameters ($\alpha_{\mathrm{T}}, N_{\mathrm{male}}$), the point and the error bars respectively represent the mean value of the statistic and its 95% confidence intervals. The x-axis is plotted on a logarithmic scale. The carrying capacity in the transition deme is calculated as: $K_{\mathrm{T}} = \frac{2.K_1.\alpha_{\mathrm{T}}}{1-\alpha_{\mathrm{T}}}$. Therefore the increase in the relative size of the transition deme $\alpha_{\mathrm{T}}$ increases the carrying capacity of the transition deme $K_{\mathrm{T}}$, but it does not affect the carrying capacities of the main demes $K_1$ and $K_2$. The overall carrying capacity $K$ is not constant in this analysis, but is higher for higher $\alpha_{\mathrm{T}}$ values. This leads to qualitatively the same results as with a constant total carrying capacity (Fig. 2). $\sigma_{\mathrm{s}} = 0.4$.

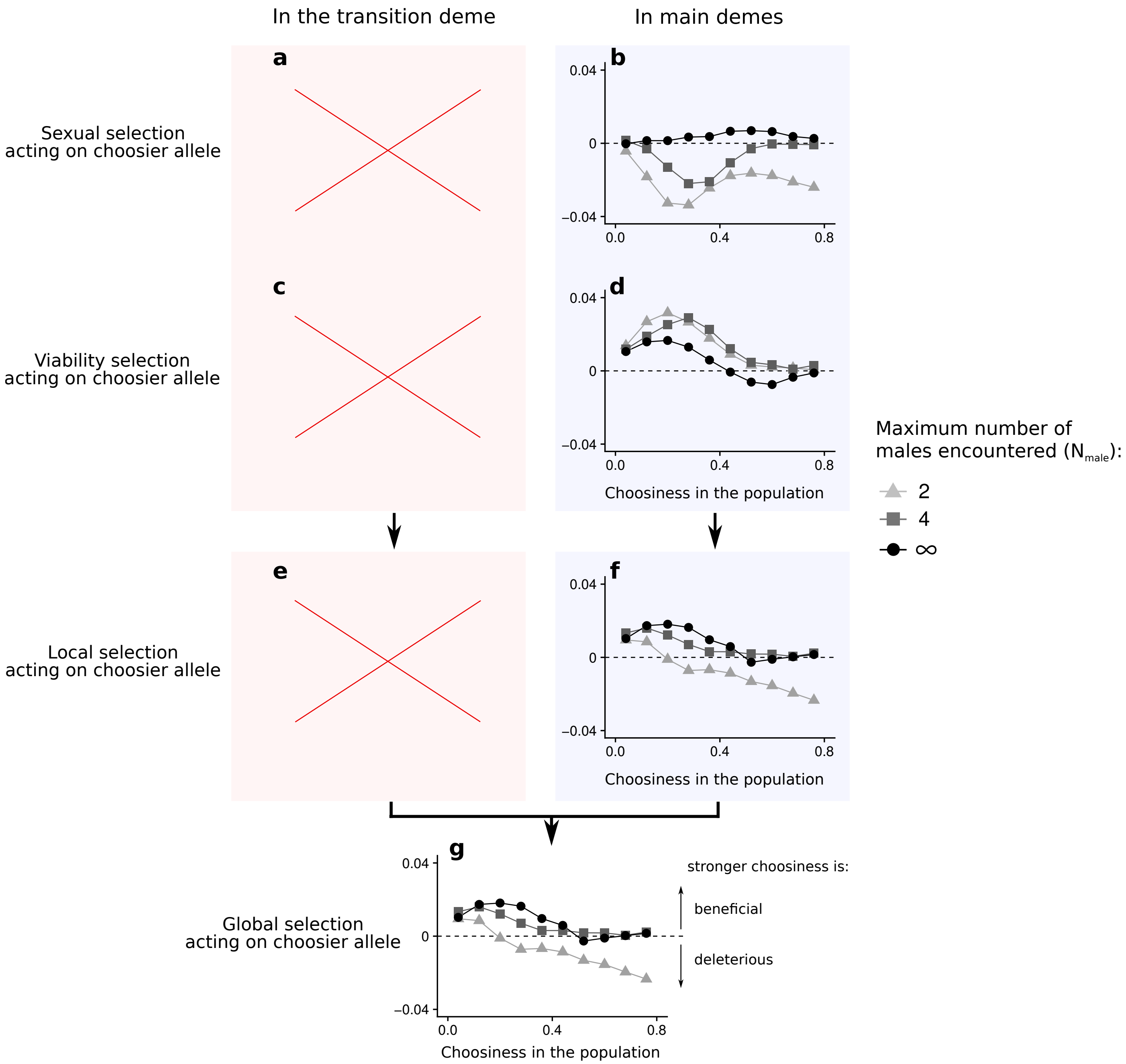


Figure S8: Local selective forces acting on choosiness in the transition deme (a, c, e) and in main demes (b, d, f), and global selection acting on choosiness (g) **when there is no transition deme, i.e., for** $\boldsymbol{\alpha_{\mathrm{T}} = 0}$. We here consider the case with $N_{\mathrm{male}} = 4$ as it corresponds to the parameter value leading to the highest choosiness at evolutionary equilibrium (Fig. 2). After the introduction of 10% of mutants with a choosier allele, local selection gradients are measured as changes in frequencies of these choosier alleles over one generation (e-f). Sexual selection (resp. viability selection) acting on alleles coding for stronger choosiness is measured as their change in frequencies during the reproduction phase (resp. viability selection phase) (a-d). Following Wickman et al. (2017), we estimate the global selection gradient as a weighted average of local selection gradients (g). $\sigma_{\mathrm{s}} = 0.4$.

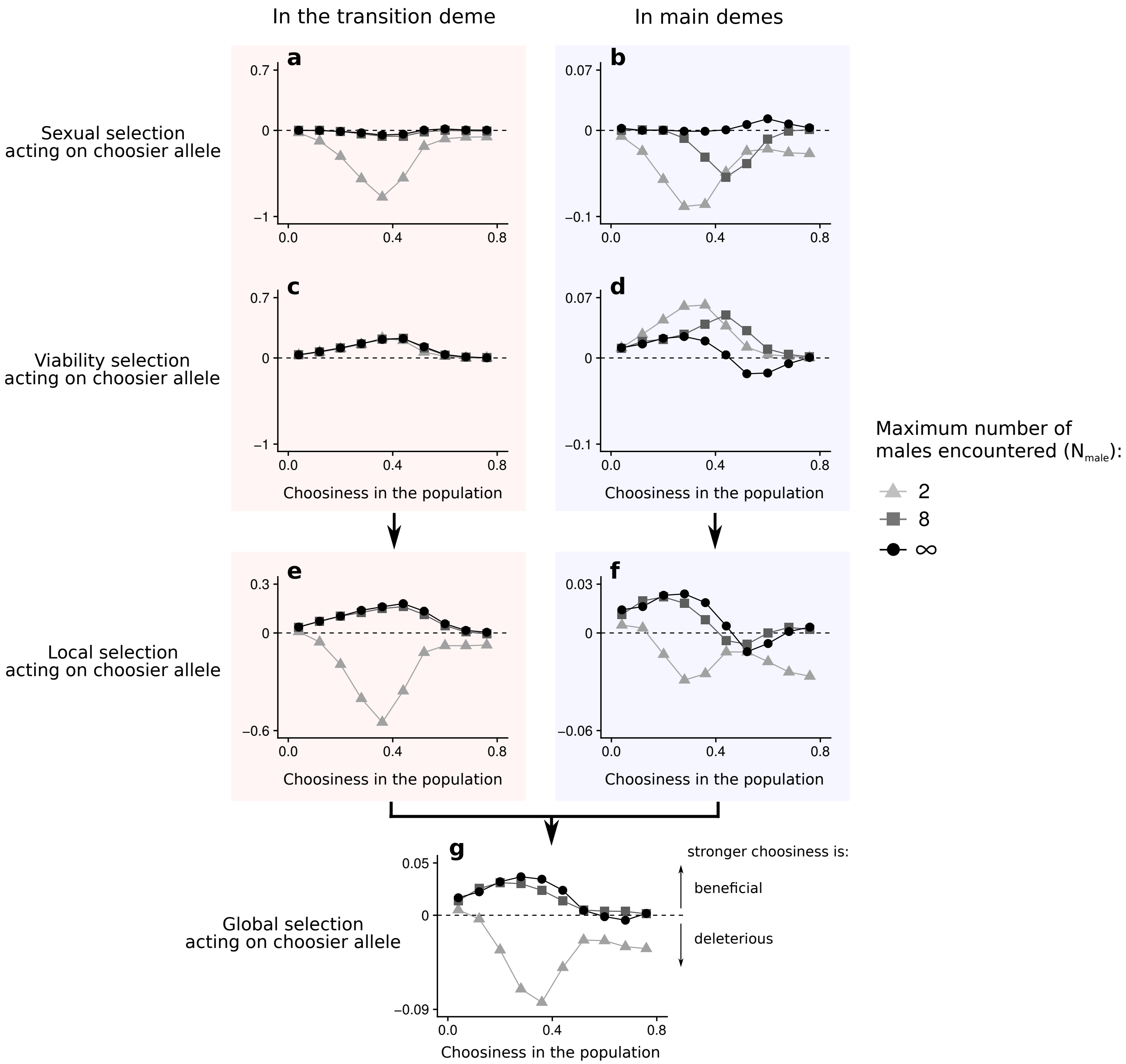


Figure S9: Local selective forces acting on choosiness in the transition deme (a, c, e) and in main demes (b, d, f), and global selection acting on choosiness (g) **for** $\boldsymbol{\alpha_{\mathrm{T}} = 0.2}$. After the introduction of 10% of mutants with a choosier allele, local selection gradients are measured as changes in frequencies of these choosier alleles over one generation (e-f). Sexual selection (resp. viability selection) acting on alleles coding for stronger choosiness is measured as their change in frequencies during the reproduction phase (resp. viability selection phase) (a-d). Following Wickman et al. (2017), we estimate the global selection gradient as a weighted average of local selection gradients (g). $\sigma_{\mathrm{s}} = 0.4$.
