## Appendix A for "Costs of choosiness can promote reproductive isolation in parapatry"

### Appendix A:
### Two-deme population genetic model

We here implement a search cost in a simple two-deme population genetic model of assortative mating via female choice (Servedio 2011). We show that a search cost (1) generates a direct selection pressure inhibiting the evolution of choosiness (because only choosy females pay the search cost), and (2) increases the nonrandom association between the allele coding for strong choosiness and alleles that are locally adapted, thereby causing indirect selection favouring strong choosiness. Here, the former effect strongly inhibits the evolution of choosiness. In our individual-based model with three demes, however, a cost of choosiness can favour the evolution of strong choosiness when selection in the transition deme compensate the direct selection pressure against choosiness caused by the cost at low levels of choosiness. In that case, the cost of choosiness causes indirect selection favouring stronger choosiness at high levels of choosiness.

#### Population genetic model

Following Servedio (2011), in this haploid model, individuals in two populations differ in the allele present at a trait locus, T; members of population 1 have predominantly trait allele $T_1$ and members of population 2 have predominantly trait allele $T_2$. The two populations exchange migrants at a rate $m$. Females are assumed to have an established mating preference for males that share their allele at this trait locus. Specifically females are $1 + \alpha_1$ times more likely to mate with a male that they prefer if they encounter one of each type of male. Strict polygyny is assumed. We incorporate selection on the trait locus T under the assumption that the more common allele in each population is favoured by selection. Selection on the trait is thus modelled by assigning fitness $1 + s$ to the trait $T_1$ in population 1 and to the trait $T_2$ in population 2 in both males and females.

Following Otto et al. (2008), we implement a search cost. Such search cost is typically called 'relative cost' due to its frequency dependence nature; even a very picky female suffers no loss in fertility if every male encountered is similar. If a female rejects a male, she may or may not be able to recuperate the lost mating opportunity. To account for this potential cost, we assume that a fraction of the time, $1 - c_{\mathrm{r}}$ (with $c_{\mathrm{r}} \in [0, 1]$), a female is able to recover the fitness lost by rejecting a dissimilar mate, and otherwise she suffers a loss in fitness.

We assume that a second locus, A, allows for the evolution of assortative mating. Specifically, alleles $A_1$ and $A_2$ at this locus code for different strengths of assortative mating, $\alpha_1$ and $\alpha_2$.

#### A search cost inhibits the evolution of choosiness

We investigate the invasion of mutant alleles $A_2$ coding for choosier mate preferences ($\alpha_2 = \alpha_1 + 1$), starting at $\alpha_1 = 0$ and assuming that the frequency of $A_2$ is equal to 0.1 initially. If allele $A_2$ increases in frequency, we do the same analysis, but this time with the resident population characterized by a choosiness $\alpha_2$ and a new mutant allele coding for even choosier mate preference ($\alpha_2 + 1$), until a choosier mutant alleles cannot incade. This choosiness value corresponds to the CSS choosiness; an allele coding for this choosiness value can always invade and cannot be replaced.

Using those numerical iterations, we can show that without a relative cost, the CSS choosiness corresponds to the choosiness value that maximize divergence at the T locus (red vertical line for $c_{\mathrm{r}} = 0$ in Figure A1) (as shown in Servedio 2011). With a relative cost, choosiness can be deleterious leading to a CSS choosiness equal to 0 (not shown in Figure A1 due to the use of log scale).

This effect is caused by a direct selection pressure inhibing choosiness: the search cost is paid by choosy females that reject unpreferred males.

#### A search cost leads to nonrandom associations between alleles coding for strong choosiness and adapted alleles

Without a search cost, maximum divergence and the CSS choosiness occurs when choosier allele $A_2$ becomes associated with the allele under negative selection locally (with $T_1$ in deme 2, i.e. $LD = \text{freq}_{A_2T_2}\text{freq}_{A_1T_1} - \text{freq}_{A_2T_1}\text{freq}_{A_1T_2} < 0$; with $T_2$ in deme 1, i.e. $LD > 0$), i.e., there is indirect selection against strong choosiness (for $c_\mathrm{r} = 0$ in Figure A1).

A search cost increases divergence for high $\alpha_1$ because rare females are not obtaining mates, so rare males have low mating success (so sexual selection, which is lost at high $\alpha_1$ when $c_\mathrm{r} = 0$, is re-established). As a result, as a search cost increases, the association between the choosier allele $A_2$ and the locally adapted allele (and therefore indirect selection favouring stronger choosiness) occurs at stronger levels of choosiness (in Figure A1, the sign of normalized LD changes at higher levels of choosiness as the relative cost increases). This means that if the cost of choosiness was not associated with a direct cost (lost mating opportunities), stronger choosiness would evolve. This is why in the three-deme model in the main text, a cost of choosiness can favour the evolution of strong choosiness when selection in the transition deme compensate the direct selection pressure against choosiness caused by the cost at low levels of choosiness.

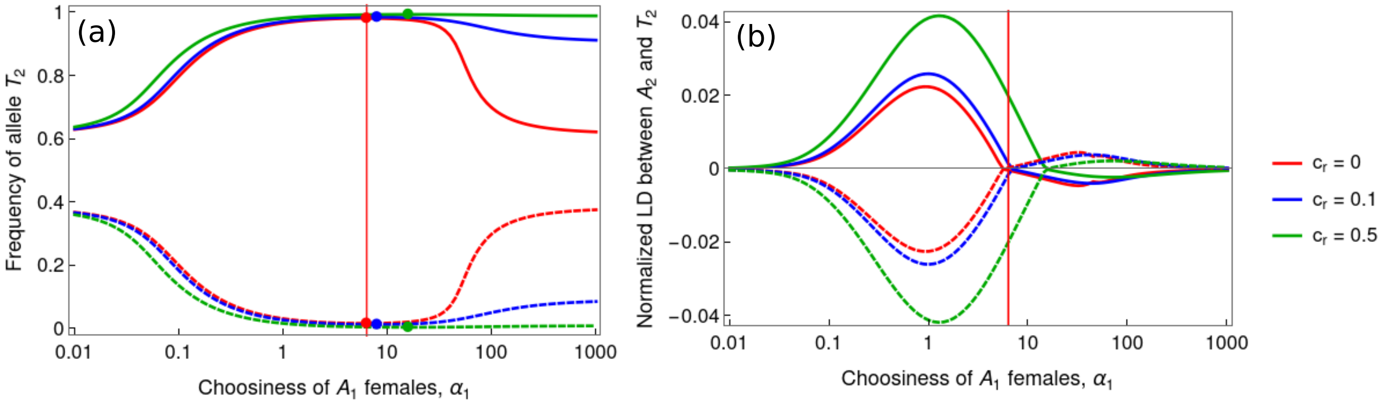


Figure A1: (a) Frequencies of $T_2$ depending on the strength of choosiness ($\alpha_1$) and on the search cost ($c_\mathrm{r}$), in deme 2 where $T_2$ is favoured (plain lines) and in deme 1 where $T_2$ is disfavoured (dashed lines). Points are drawn at the choosiness value maximum divergence, and the red line represents the CSS choosiness obtained for $c_\mathrm{r} = 0$ (CSS choosiness is 0 for $c_\mathrm{r} = 0.1$ and $c_\mathrm{r} = 0.5$, not shown in the graph). Here we represent the equilibrium state when all individuals carry allele $A_1$ (i.e., when there is no mutant; obtained numerically). (b) Normalized linkage disequilibrium (LD) between alleles $A_2$ and $T_2$ in deme 2 (plain lines) and in deme 1 (dashed lines) assessed 3 generations after the inclusion of the mutant allele at frequency 0.1. We normalize LD by dividing it by the theoretical maximum difference between the observed and expected haplotype frequencies. As a search cost increases, the association between the choosier allele $A_2$ and the locally adapted allele (and therefore indirect selection favouring choosiness) occurs at stronger levels of choosiness. This means that if the search cost was not associated with a direct cost (lost mating opportunities), stronger choosiness would evolve. Parameter values: $s = 0.01$, $m = 0.01$
